# Long-term decline in egg size of Japanese hatchery chum salmon links to declines in return rate and abundance

**DOI:** 10.64898/2026.05.30.729012

**Authors:** Shuichi Kitada

## Abstract

Variation in life-history traits contributes to fitness and population productivity in salmonids. To investigate long-term changes in egg size and body size of Japanese hatchery chum salmon and their links to declines in return rate and abundance, I analyzed egg size and female fork length (FL) of approximately 21,000 age-4 and age-5 fish from 13 hatchery-enhanced rivers in Japan during 1999–2019. River-specific generalized additive models (GAMs) identified synchronous declines in egg size across most rivers. Hierarchical GAMs adjusting for female FL, age, and river effects revealed a nationwide decline in egg size, whereas adjusted FL showed pronounced interannual variation without a long-term decline. Mean age at return also declined after approximately 2010 because of shifts in age composition. Standardized exponential decay rates were remarkably similar for egg size, return rate, and chum salmon abundance, suggesting common underlying environmental and evolutionary processes. These findings suggest that long-term declines in maternal investment may have contributed to declining return rates and abundance, underscoring the importance of life-history traits in evaluating the long-term consequences of hatchery enhancement programs.

## 1. Introduction

Variation in life-history traits contributes to fitness and population productivity in salmonids (Quinn 2005; Koch and Narum 2021). Among these traits, female body size is a key reproductive characteristic because larger females generally produce larger eggs. Larger eggs generally produce larger offspring with higher early survival and growth, effects that can carry over to later life stages and influence recruitment, thereby playing a central role in lifetime reproductive success and reflecting maternal investment strategies that balance fecundity, offspring quality, and environmental uncertainty (Beacham and Murray 1985, 1993; Fleming and Gross 1990; Einum and Fleming 1999, 2000; Heath et al. 1999). Consequently, variation in female body size and egg size is closely linked to fitness and population productivity (Chambers and Leggett 1996; Mousseau and Fox 1998; Quinn et al. 1995).

Over recent decades, changes in reproductive life-history traits have been increasingly documented in Pacific salmon (*Oncorhynchus* spp.) populations. Among these, declines in body size have been widely reported and are considered to have important consequences for population productivity and resilience. These declines are attributed largely to shifts in age structure driven by climate change and increasing competition at sea (Lewis et al. 2015; Jeffrey et al. 2017; Oke et al. 2020; Ohlberger et al. 2023). Similar long-term declines in body size have also been reported in Atlantic salmon (*Salmo salar*) populations (Fiske et al. 2025). Among reproductive traits, reductions in egg size have been reported in heavily supplemented Pacific salmon populations (Heath et al. 2003), whereas other studies found no consistent differences in egg size between hatchery and wild fish (Beacham and Murray 1993; Quinn et al. 2004; Beacham 2010). Declines in fecundity have also been observed in hatchery-reared Chinook salmon (*O. tshawytscha*) populations (Malick et al. 2023).

Hatchery propagation has long been used to enhance marine and anadromous fish populations and support fisheries since the mid-19th century (Svåsand et al. 2000; Naish et al. 2007). Each year, more than 2.6 × 10^10^ juveniles of approximately 180 marine species, including salmon, are released into the wild in over 20 countries (Kitada et al. 2018), making hatchery enhancement one of the world’s most widely applied fisheries management and conservation strategies

For Pacific salmon, hatchery enhancement programs have operated throughout the Pacific Rim, enhance fishery production, and compensate for habitat degradation, making them among the largest and most persistent human interventions in aquatic ecosystems worldwide (Naish et al. 2007; Amoroso et al. 2017; Kitada 2018). Hatchery-origin fish now constitute a substantial proportion of salmon biomass in the North Pacific (Ruggerone and Irvine 2018). In addition, hatchery propagation has become an important conservation tool for Pacific salmon and Atlantic salmon populations listed under the U.S. Endangered Species Act and for threatened populations in Canada (Maynard and Trial 2014).

Although these programs have substantially contributed to fishery yields (Hilborn and Eggers 2000), and have helped stabilize some Pacific salmon populations (Ford et al. 2025), growing evidence suggests that prolonged artificial propagation can generate complex eco-evolutionary effects, including reduced reproductive success (Araki et al. 2007; Christie et al. 2012a; Christie et al. 2014; Shedd et al. 2022), lower survival rates (Reisenbichler and Rubin 1999), declines in effective population size (Christie et al. 2012b). Additional studies have reported differential expression of numerous genes (Christie et al. 2016), widespread differences in DNA methylation (Le Luyer et al. 2017; Gavery et al. 2018), and changes in adaptive genetic variation associated with age at maturity and body size in hatchery salmonids (Harris et al. 2026). Such genetic changes associated with prolonged hatchery propagation may ultimately reduce the resilience and long-term sustainability of supplemented populations (Ryman and Laikre1991; Waples 1991; Waples and Drake 2004).

Japanese chum salmon (*O. keta*) provide a uniquely valuable system for addressing this issue because Japan operates one of the world’s largest stock enhancement programs for this species (Amoroso et al. 2017; Kitada 2018). Because the feasibility and risks of hatchery enhancement can only be rigorously evaluated at large spatial and temporal scales (Hilborn 2004), Japanese chum salmon offers a rare opportunity to assess the long-term biological consequences of artificial propagation programs. Despite continued large-scale releases, adult returns have shown a marked long-term decline since the early 2000s (Fig. S1), accompanied by reductions in return rate (Kitada et al. 2025). Although geographic variation and temporal trends in egg size have also been reported, previous studies relied on incomplete temporal coverage across rivers and years (Hasegawa et al. 2021; Kitada et al. 2025). Long-term changes in the age structure of Japanese chum salmon remain poorly understood.

In this study, I examined long-term trends in body size and egg size of approximately 21,000 female chum salmon from 13 hatchery-enhanced rivers in Japan over 21 years (1999–2019). I first described spatial variation in female fork length (FL) and egg size among rivers to provide a baseline for interpreting temporal changes across populations, then characterized temporal trends using generalized additive models (GAMs) fitted to individual-level data, and finally estimated nationwide trends using hierarchical GAMs, with egg-size trends adjusted for female FL, age, and river. To assess temporal changes in age at maturity, I also analyzed annual age composition and mean age at return using age-composition data from approximately 1.2 × 10^6^ returning adults collected during 1997–2020. Together, these analyses provide new insights into long-term changes in key life-history traits and their potential links to declining return rates and population abundance.

## 2. Methods

### 2.1 Data

I used adult chum salmon trait data compiled in Supplementary Data S3 of Kitada et al. (2025), which were originally obtained from the Fisheries Research and Education Agency Salmon Database (FRA 2026). The available dataset included of female fork length (FL) and egg size measurements from age-4 and age-5 chum salmon collected from 13 hatchery-enhanced rivers in Japan, with observations spanning 1997–2020 (5 rivers in Hokkaido and 8 rivers in Honshu; *n* = 14,084 fish for age-4 and *n* = 9,583 fish for age-5) (Fig. 1A, Table S1).

**Fig. 1.**
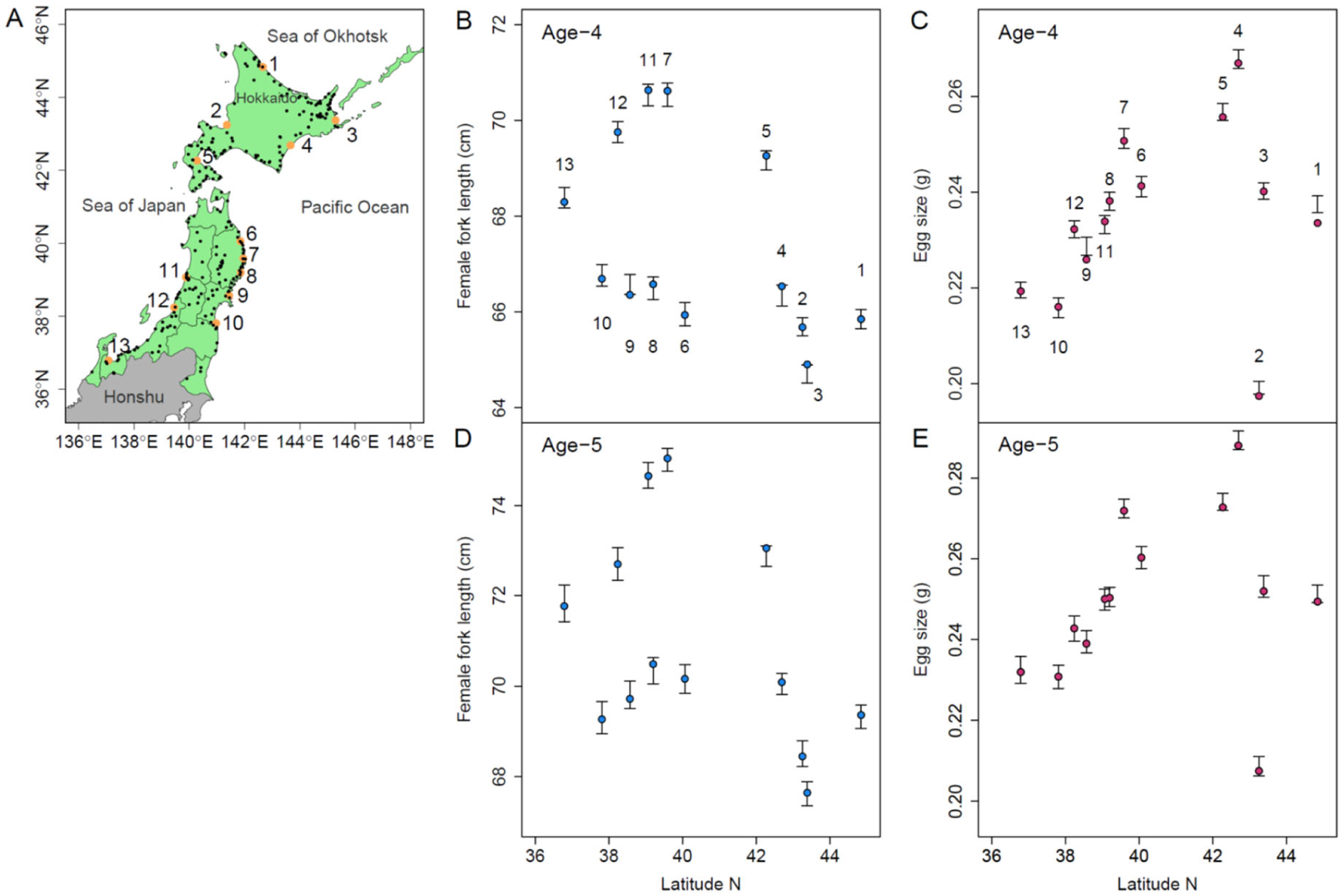
Chum salmon were sampled from 13 hatchery-enhanced rivers in Japan. (A) Orange dots indicate the mouths of the sampled rivers, while the chum salmon return area is shown in green, with salmon hatcheries indicated by black dots (adapted from Kitada, 2000). Geographic distributions of mean female fork length (FL) for age-4 (B) and age-5 (D), and mean egg size for age-4 (C) and age-5 (E) hatchery-origin chum salmon sampled during 1999–2019 (*n* = 12,428 for age-4 and *n* = 8,862 for age-5). Blue symbols in panels B and D represent river-specific mean FL, and red symbols in panels C and E represent river-specific mean egg size; error bars indicate 95% confidence intervals of the river-specific means calculated from all screened individual observations within each river during the study period.

These rivers were selected because they provided the longest and most complete time series available for female FL and egg size (Tables S2 and S3). Ages were determined by scale analysis (Hasegawa et al. 2021). Chum salmon are semelparous, with adults returning once to spawn before dying. Returning adults were predominantly age-4 and age-5 fish, although the proportions of each age class varied among rivers.

For analyses of age composition and mean age at return, I used age-composition data for returning adults during 1997–2020 (Supplementary Data S2 in Kitada et al. 2025, FRA 2026), comprising 1,210,717 fish collected from 1,902 surveys in 52 Hokkaido rivers and 102 Honshu rivers. The data consisted of age-2 to age-8 fish, and age-specific numbers were estimated from the reported age composition and sample size for each survey.

I compiled chum salmon catch and hatchery release statistics from the North Pacific Anadromous Fish Commission (NPAFC 2025; 1952–2024) and prefecture-level statistics for Japan from FRA (FRA 2026; 1970–2025). Long-term historical records of chum salmon hatchery releases and catches in Japan (1870–2025) were updated from Kitada (2020). Japanese chum salmon fisheries have been based largely on the capture of returning fish in coastal waters, including rivers, with limited escapement to natural spawning populations. The high-seas salmon fishery has been closed since 1992 (Morita et al. 2006). Therefore, catch was used as an index of annual chum salmon abundance in this study.

### 2.2 Within-year variation and data screening

Individual measurements of female FL and egg size showed substantial variation within years across all rivers (Fig. S2–S5). A total of 10 biologically implausible observations were excluded before analysis based on biological plausibility, including six age-4 females with egg size ≥ 0.4 g, two age-4 females with FL ≥ 85 cm, and two age-5 females with FL ≤ 50 cm. Although these datasets covered most of the 1997–2020 period, observations were unavailable for four rivers in 1997 and 1998 and for three rivers in 2020. Therefore, all subsequent analyses were restricted to 1999–2019, during which data were available for nearly all river-year combinations, with only seven river-years missing across the 13 rivers. The resulting dataset, comprising 21,290 females (12,428 age-4 and 8,862 age-5 females), was used in all subsequent analyses.

### 2.3 River-level variation in female body size and egg size

To describe spatial variation in female FL and egg size, I calculated river-specific mean values and their 95% confidence intervals using all individuals sampled during the study period. River-specific means were plotted against river-mouth latitude separately for age-4 and age-5 females to illustrate geographic patterns.

### 2.4 River-specific temporal trends in female body size and egg size

To characterize temporal patterns in female FL and egg size across the 13 rivers, I fitted generalized additive models (GAMs) using the mgcv package in R version 4.5.2 (Wood 2017; R Core Team 2025).

The models were specified as

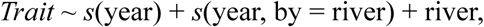

where *Trait* represents female FL or egg size. Gaussian errors were assumed, and models were fitted using restricted maximum likelihood (REML). The smooth term *s*(year) describes the overall temporal trend, whereas *s*(year, by = river) allows river-specific deviations from the overall trend. River was included as a categorical factor to account for differences in mean trait values among rivers. Model adequacy was assessed using residual diagnostic plots and the gam.check() function.

For visualization of the global temporal pattern, a separate GAM, *Trait* ∼ *s*(year), with a single common smooth of year was fitted to all observations pooled across the 13 rivers. The resulting smooth was presented as the global smooth in Figures 2 and 3 and Figures S6 and S7.

**Fig. 2.**
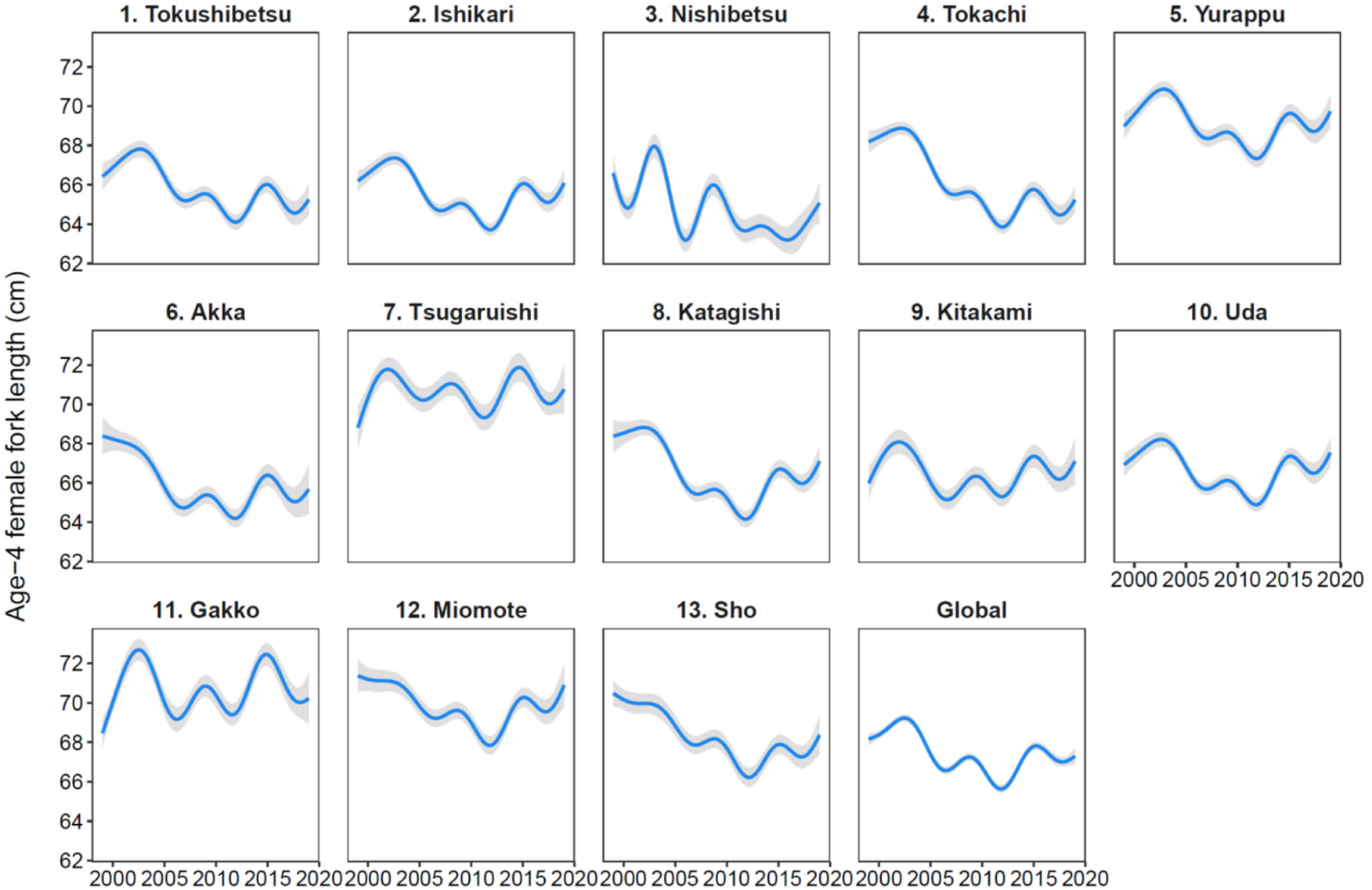
Predicted temporal trends in female fork length (FL) for age-4 chum salmon estimated from generalized additive models fitted to individual-level data from 12,428 females sampled from 13 hatchery-enhanced rivers for the 1999–2019 brood years after excluding 8 outliers. Blue lines show the fitted GAM smooths, and gray shaded areas indicate the 95% confidence intervals. The global smooth represents the common temporal trend estimated from a separate GAM with a single common smooth of year fitted to all observations pooled across the 13 rivers.

**Fig. 3.**
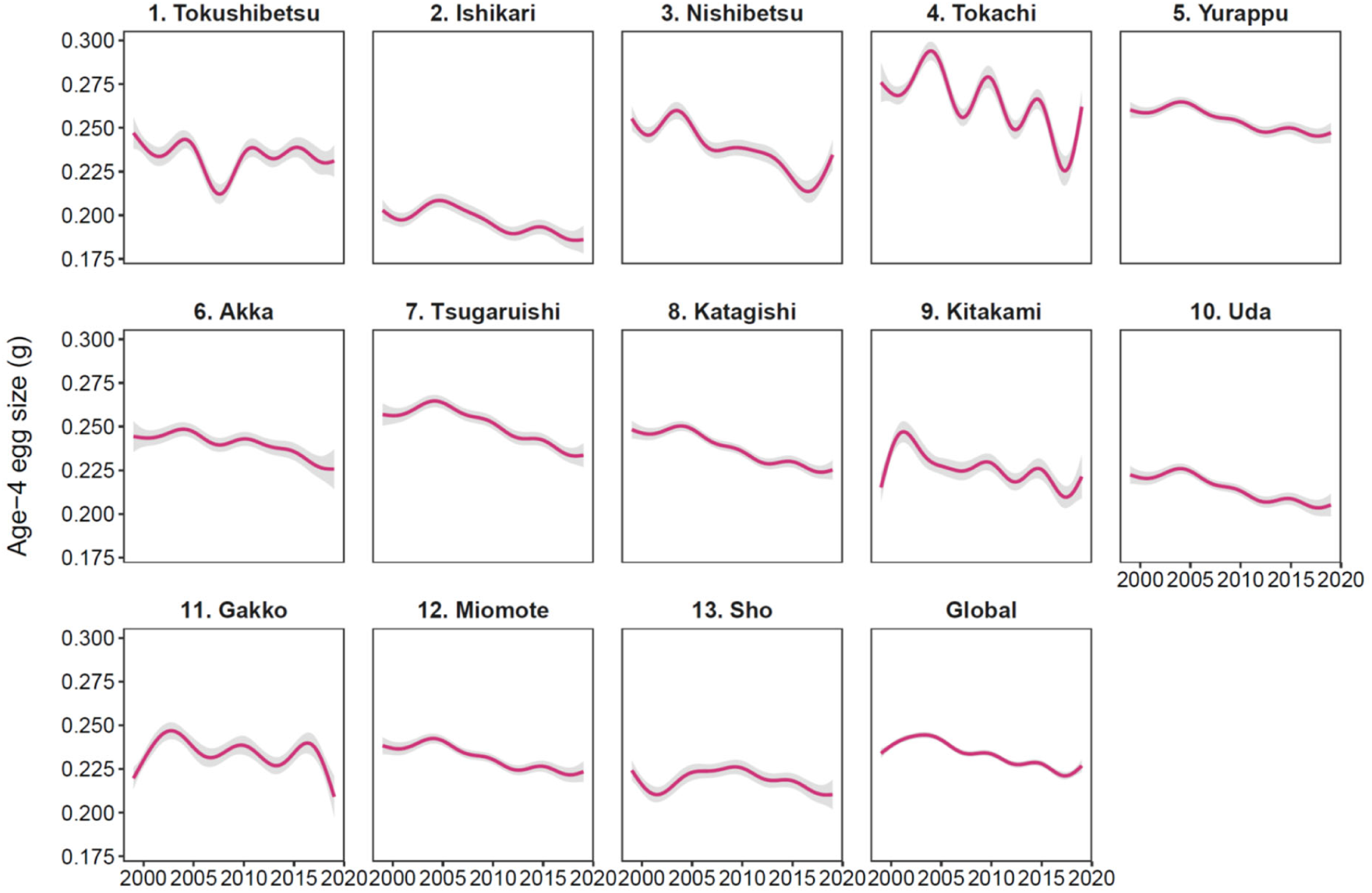
Predicted temporal trends in egg size for age-4 chum salmon estimated from generalized additive models fitted to individual-level data from 12,428 females sampled from 13 hatchery-enhanced rivers for the 1999–2019 brood years after excluding 8 outliers. Pink lines show the fitted GAM smooths, and gray shaded areas indicate the 95% confidence intervals. The global smooth represents the common temporal trend estimated from a separate GAM with a single common smooth of year fitted to all observations pooled across the 13 rivers.

The default basis dimension (*k* = 10) was used for all smooth terms. Age and female FL were not included as covariates because the objective of this analysis was to visualize observed temporal trajectories within each river rather than estimate trends adjusted for individual-level characteristics. The potential effects of these individual-level characteristics on temporal trends were examined separately in the subsequent analysis (See Section 2.5).

### 2.5 Nationwide temporal trends in female body size and egg size

To estimate nationwide temporal trends in egg size and female fork length (FL), I fitted hierarchical GAMs (Pedersen et al. 2019) to individual-level data using the mgcv package.

The FL model was specified as

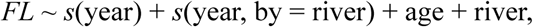

where *s*(year) represents the overall temporal trend and *s*(year, by = river) allows river-specific deviations from the overall trend. Age and river were included as categorical fixed effects.

The egg-size model was specified as

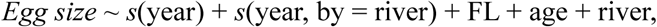

where *s*(year) represents the overall temporal trend and *s*(year, by = river) allows river-specific deviations from the overall trend. Female FL was included as a continuous covariate, whereas age and river were included as categorical fixed effects. Smoothness parameters were estimated using REML, using the default basis dimension (*k* = 10) for all smooth terms.

Alternative GAM structures were compared using Akaike’s Information Criterion (AIC, Akaike 1973). Candidate models included a simpler GAM with a single global smooth for year and the hierarchical GAM described above, which included additional river-specific smooths for year. To evaluate the effect of basis dimension, the hierarchical GAM was also fitted with restricted basis dimensions (*k* = 6) for both smooth terms. The model with the lowest AIC was selected for subsequent analyses. Model adequacy was assessed using residual diagnostic plots and the gam.check() function.

### 2.6 Changes in age composition and mean age at return

Age composition data of returning adults (1997–2020) were used to estimate annual age composition and mean age at return (*N* = 1,210,717 fish), using the longest available time series to characterize long-term changes in age structure. The weighted mean proportion of age-*a* fish in year *t* was calculated as 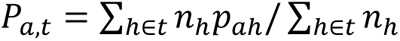, where ℎ denotes each survey (*h* = 1, …, 1,902) in 154 rivers in Hokkaido and Honshu, *p_a_*_ℎ_ is the observed proportion of age-*a* fish in survey ℎ, *n*_ℎ_ is the sample size of survey ℎ, and *a* is age at return (*a* = 2, …,8). The annual weighted mean age at return was then calculated as 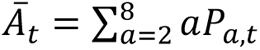.

Changes in mean age at return were evaluated using GAM, with annual weighted mean age at return modeled as a smooth function of year: *Ā_t_* ∼ *s*(year). The model was fitted using restricted maximum likelihood (REML), using the default basis dimension (*k* = 10) for the smooth term.

The overall weighted mean age at return across all sampled fish during 1997–2020 was calculated as 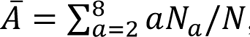, where *N_a_* is the total number of sampled fish of age *a* across all surveys conducted during 1997–2020. The weighted mean age at return across all sampled fish (*Ā*) was used as the generation time for estimating exponential decay rates per generation in life-history traits and population dynamics (See Section 2.7).

### 2.7 Parallel declines in life-history traits and population dynamics

I quantified temporal declines in age-4 and age-5 egg size, brood-year-specific return rates, and abundance of Japanese chum salmon using an exponential decay model, *N* = *ae*^−*bx*^, where *x* represents time (years), *a* is the intercept, and *b* is the instantaneous decay rate. Brood-year-specific return rates were calculated as the cumulative number of returning adults from ages 2 to 5 divided by the number of juveniles released in each brood year (1999–2019) (Kitada et al. 2025). Exponential decay models were fitted to age-4 egg size, age-5 egg size, return rate, and return abundance. Model assumptions were evaluated using standard linear model diagnostic plots, including residuals versus fitted values, normal Q–Q plots, scale-location plots, and residual leverage plots. Mean age at return was analyzed separately and was not included in the exponential decay analysis.

Exponential decay models were used because they assume a constant instantaneous rate of decline over time and allow comparison of proportional rates of change among biological variables. Such models have been widely applied in fisheries science and conservation (Beverton and Holt 1957; Hilborn and Mangel 2013). Araki et al. (2007) fitted an exponential decay model to published estimates of relative reproductive success (RRS) across generations for captive-reared salmonids. RRS was defined as the number of adult offspring produced per hatchery-origin parent relative to that produced per wild-origin parent. Following this approach, annual instantaneous decay rates (*b̂*) estimated in the present study were converted to per-generation rates using the generation time estimated from the mean age at return (*Ā*) and compared with the per-generation rate estimated by Araki et al. (2007). Egg size, return rate, and abundance were log-transformed and then standardized to enable comparisons among variables measured on different scales. Instantaneous decay rates were estimated from the fitted slopes, and models with and without a breakpoint were compared using AIC, with segmented regression models fitted using the segmented package in R.

Finally, I examined relationships among female FL, egg size, and return rate using ordinary least-squares linear regression. The relationship between mean female FL and mean egg size was examined using combined age-4 and age-5 observations for brood years 1999–2018 (20 observations for each age class; *n* = 40). The relationship between mean egg size and return rate was then tested using mean egg size across age-4 and age-5 fish for brood years 1999–2018 and return rates for the corresponding release years 2000–2019 (*n* = 20), with return rates for hatchery-reared chum salmon (ages 2–5) reported by Kitada et al. (2025).

## 3. Results

### 3.1 River-level variation in female body size and egg size

Female FL varied substantially among river populations in both age classes (Fig. 1B, D). The largest females were observed in the Gakko River population (#11; Honshu Sea of Japan coast) and the Tsugaruishi River population (#7; Honshu Pacific coast), whereas the smallest females occurred in the Nishibetsu River population (#3; Nemuro Strait) (Fig. 1A).

Egg size also showed pronounced variation among rivers (Fig. 1C, E). The largest eggs were observed in the Tokachi River population (#4; Hokkaido Pacific coast), whereas the smallest occurred in the Ishikari River population (#2; Hokkaido Sea of Japan coast). Geographic patterns in FL and egg size were generally consistent between age-4 and age-5 females, respectively.

### 3.2 River-specific temporal trends in female body size and egg size

Generalized additive models (GAMs) fitted to individual-level data showed that individual age-4 FL measurements were widely scattered around the fitted smooths in all rivers (Figure not shown). Despite this substantial within-river variation, most populations exhibited only modest temporal fluctuations in female FL. Gradual declines were evident only in a few populations, particularly the Tokachi, Akka, Katagishi, and Sho rivers.

In contrast, individual age-4 egg-size measurements also showed considerable within-river variation, but GAM smooths revealed a consistent long-term decline in most river populations (Fig. 3). Although the magnitude and timing of the decline varied among rivers, downward trends were broadly synchronous across populations. Similar temporal patterns were observed for age-5 fish (Figs. S6 and S7).

The river-specific GAMs explained 31.3–38.0% of the deviance across the four models for female FL and egg size in age-4 and age-5 fish (adjusted *R*^2^ = 0.31–0.38). Residual diagnostic plots for the river-specific GAMs for age-4 and age-5 female FL and egg size did not indicate major violations of model assumptions (Figs. S8–S11). Detailed results of the river-specific GAMs, including parametric coefficients, edf, and *p*-values for the smooth terms, are provided in Tables S4–S11.

The global smooths, representing the common temporal trends estimated from all observations pooled across the 13 rivers, were consistent with the river-specific patterns for both age-4 and age-5 female FL and egg size (Figs. 2, 3, S6, and S7, bottom-right panels). The global smooths explained 4.0–8.9% of the deviance across the four models for female FL and egg size in age-4 and age-5 fish (adjusted *R*² = 0.04–0.09). For age-4 fish, the global smooths were highly significant for female FL (effective degrees of freedom [edf] = 8.78, *F* = 89.4) and egg size (edf = 8.23, *F* = 55.95), with similarly significant temporal patterns for age-5 fish (edf = 8.85, *F* = 95.02 for FL; edf = 8.64, *F* = 53.76 for egg size; *p* < 0.0001 for all models).

**Table 1.** Instantaneous annual decay rates of egg size, return rate, and abundance in Japanese chum salmon, estimated from raw and standardized data. Exponential decay models, *N* = *ae*^−*bx*^, were fitted to egg size and return rate, whereas a segmented regression was fitted to abundance (see Fig. 6).

| Data | Dependent variable | $\hat{b}$ | se | $t$ | $p$ | Adjusted $R^2$ |
| --- | --- | --- | --- | --- | --- | --- |
| Raw | Age-4 egg size | 0.0045 | 0.0008 | -5.33 | < 0.0001 | 0.58 |
|  | Age-5 egg size | 0.0056 | 0.0009 | -6.25 | < 0.0001 | 0.66 |
|  | Return rate | 0.0606 | 0.0103 | -5.87 | < 0.0001 | 0.63 |
|  | Abundance, before 2005.3 | -0.0705 | 0.0450 | 1.57 | 0.1310 |  |
|  | Abundance, after 2005.3 | 0.0862 | 0.0092 | -9.34 | < 0.0001 | 0.83 |
| Standardized | Age-4 egg size | 0.1248 | 0.0234 | -5.33 | < 0.0001 | 0.58 |
|  | Age-5 egg size | 0.1321 | 0.0212 | -6.25 | < 0.0001 | 0.66 |
|  | Return rate | 0.1294 | 0.0221 | -5.87 | < 0.0001 | 0.63 |
|  | Abundance, before 2005.3 | -0.1231 | 0.0785 | 1.57 | 0.1310 |  |
|  | Abundance, after 2005.3 | 0.1505 | 0.0161 | -9.34 | < 0.0001 | 0.83 |

For abundance, decay rates were estimated separately before and after the estimated breakpoint (2005.3 years) using segmented regression. Negative values of $\hat{b}$ indicate increasing trends. Standardized values were calculated after scaling each dependent variable to a mean of zero and a standard deviation of one.

### 3.3 Nationwide temporal trends in female body size and egg size

Hierarchical GAMs fitted to individual-level data from 13 rivers revealed contrasting nationwide temporal trends in female FL and egg size (Fig. 4).

**Fig. 4.**
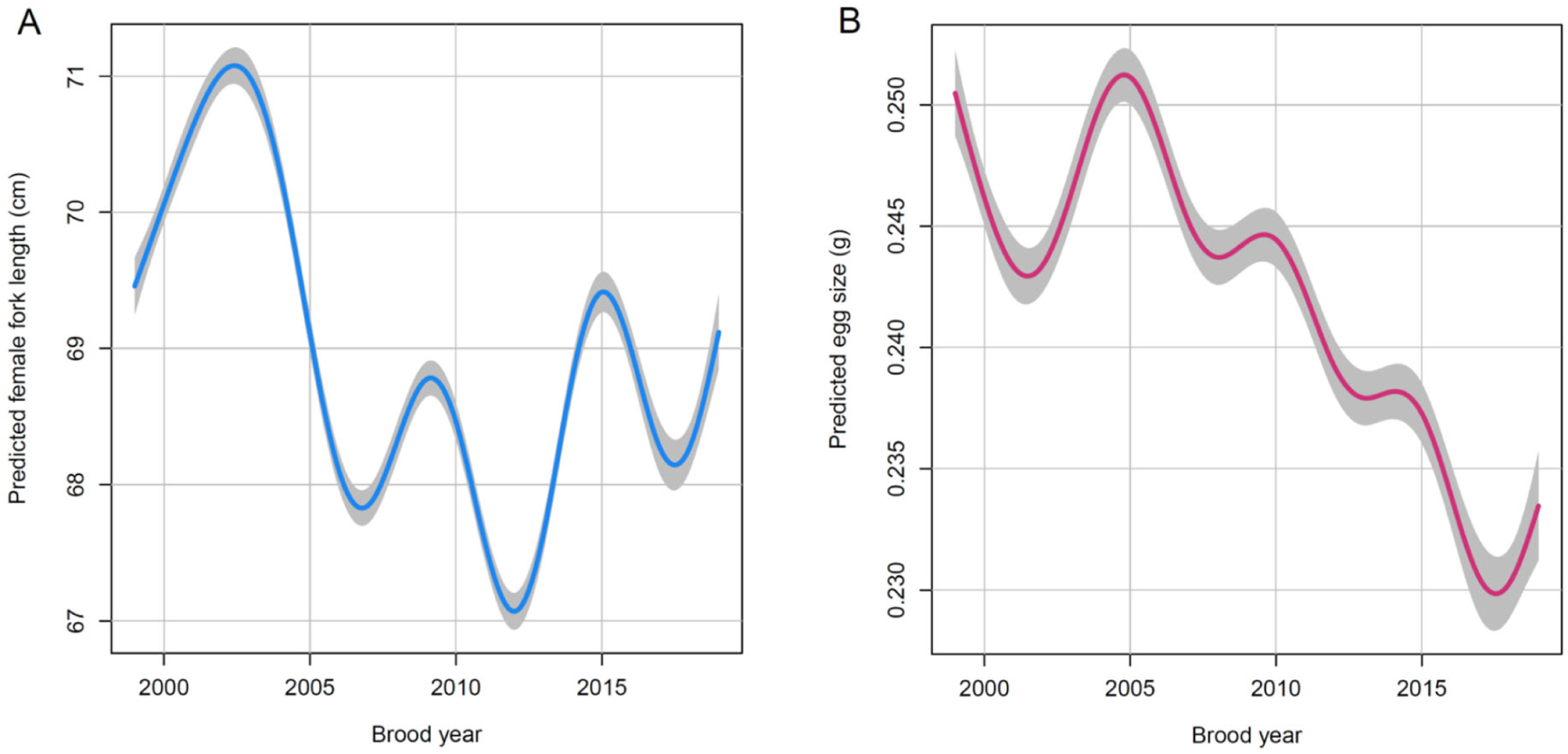
Predicted nationwide mean egg size and female fork length (FL) estimated from hierarchical generalized additive models fitted to individual-level data from 21,290 age-4 and age-5 female chum salmon sampled from 13 hatchery-enhanced rivers after excluding 10 outliers. Female FL was adjusted for age and river effects, whereas egg size was adjusted for female FL, age, and river effects. (A) Predicted female FL (adjusted *R*² = 0.44; deviance explained = 43.7%). (B) Predicted egg size (adjusted *R*² = 0.42; deviance explained = 41.9%). Blue and pink lines show the fitted GAM smooths, and the shaded areas indicate 95% confidence intervals. Complete parametric coefficients and the approximate significance of smooth terms for these models are provided in Tables S12–S15.

After accounting for age and river effects, predicted mean FL fluctuated among brood years, with peaks around 2003, 2009, and 2015, but showed no consistent long-term decline (Fig. 4A). The model explained 43.7% of the deviance (adjusted *R*^2^ = 0.44). Residual diagnostic plots did not indicate major violations of model assumptions for either hierarchical GAM (Fig. S12).

Age was positively associated with female FL (*p* < 0.0001), and the overall smooth effect of year remained highly significant after accounting for the effects of age and river (edf = 8.91, *F* = 60.11, *p* < 0.0001). River effects indicated significant spatial differences in mean female FL among all rivers (Table S12). Nine rivers showed significant river-specific smooth terms, indicating temporal trajectories that differed significantly from the nationwide trend, whereas the remaining four rivers showed no significant departures from the nationwide trend (Table S13).

For the egg-size analysis, the hierarchical GAM including river-specific smooths provided a substantially better fit than both a simpler GAM with a single global smooth for year but no river-specific smooths (ΔAIC = 409) and the hierarchical GAM with restricted basis dimensions (*k* = 6; ΔAIC = 237). Using this model, predicted mean egg size, after accounting for female FL, age, and river effects, showed a marked decline over the study period (Fig. 4B). The decline accelerated after 2005 and continued until the late 2010s. The hierarchical GAM explained 41.9% of the deviance (adjusted *R*^2^ = 0.42) for egg size. Residual diagnostic plots did not indicate major violations of model assumptions for either hierarchical GAM (Fig. S13).

Female FL and age were both positively associated with egg size (both *p* < 0.0001), and the overall smooth effect of year remained highly significant after accounting for the effects of female FL, age, and river (edf = 8.43, *F* = 17.48, *p* < 0.0001). River effects indicated significant spatial differences in mean egg size among most rivers (Table S14). Seven rivers showed significant river-specific smooth terms, indicating temporal trajectories that differed significantly from the nationwide trend, whereas the remaining six rivers showed no significant departures from the nationwide trend (Table S15).

### 3.4 Changes in age composition and mean age at return

The age composition of returning chum salmon shifted after around 2010. Despite considerable interannual variation, the proportion of age-6 and age-5 fish tended to decline, whereas the proportions of age-3 and, to a lesser extent, age-4 fish increased. Consequently, the weighted mean age at return declined markedly during the latter part of the study period (Fig. 5). The GAM fitted to weighted annual mean age at return indicated a significant nonlinear temporal trend (edf = 3.97, *F* = 8.29, *p* < 0.001; adjusted *R*^2^ = 0.63; deviance explained = 69.6%). Mean age at return increased gradually from 1997 to around 2010, followed by a rapid decline, indicating a marked shift toward younger ages at return. The weighted overall mean age at return across all returning adults included in the age-composition dataset (Ā) was estimated as 4.2 years.

**Fig. 5.**
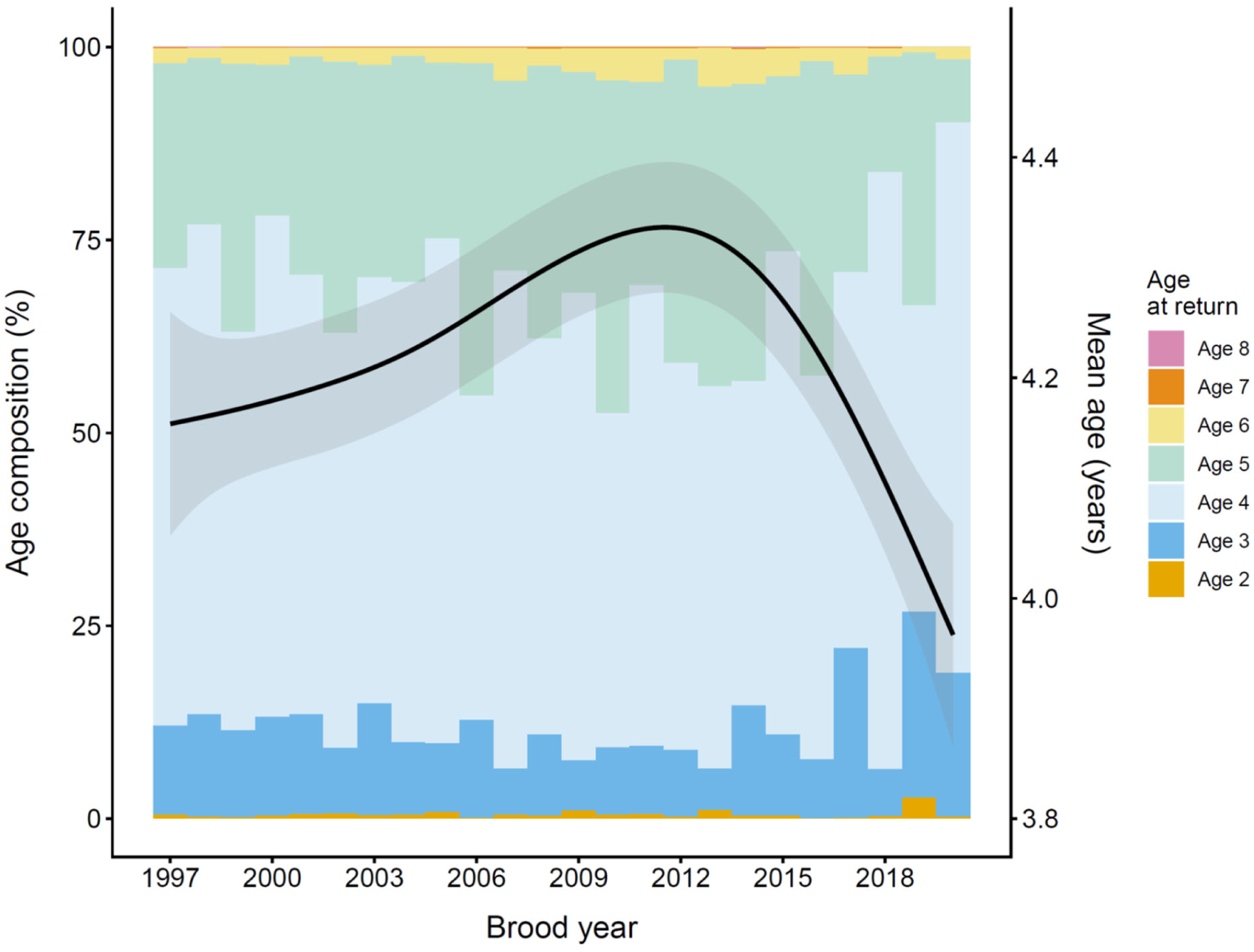
Temporal changes in the age composition of returning Japanese chum salmon based on age composition data from 1,210,717 returning adults collected from 154 rivers during 1997– 2020. The annual weighted mean age at return is overlaid with a GAM-estimated trend to illustrate the long-term pattern (edf = 3.97, *F* = 8.29, *p* < 0.001; adjusted *R*² = 0.63; deviance explained = 69.6%); the black line shows the fitted trend and the grey band its 95% confidence interval.

### 3.5 Parallel declines in life-history traits and population dynamics

Residual diagnostic plots for the linear exponential decay models showed no major violations of model assumptions for egg size and return rate (Figs. S14–S16), whereas the residuals for chum salmon abundance showed a quadratic pattern (Fig. S17), indicating a potential non-linear relationship. Segmented regression provided little improvement for age-4 egg size and return rate (ΔAIC = 0.14 and 0.33, respectively), whereas the linear model was better supported for age-5 egg size (ΔAIC = −1.65). In contrast, the segmented model was strongly supported for chum salmon abundance (ΔAIC = 12.23; Table S16). Although variation in the observed values was substantial, all annual observations for egg size, return rate, and chum salmon abundance remained within the 95% prediction intervals except for chum salmon abundance in 2022 and 2025 (Fig. 6).

The non-standardized instantaneous exponential decay rates were approximately 0.005 per year for egg size and 0.061 per year for return rate. For chum salmon abundance, the decay rate changed from −0.071 per year before the breakpoint to 0.086 per year after the breakpoint, with the breakpoint estimated at 2005.3 (Table 1), indicating a shift from an increasing to a declining trend around 2005. Although egg size declined much more slowly in absolute terms, standardized decay rates were remarkably similar among these three variables, with values of 0.125 and 0.132 for age-4 and age-5 egg size, 0.129 for return rate, and 0.151 for chum salmon abundance after the breakpoint, respectively (Table 1).

Using a generation time of 4.2 years (Ā), these non-standardized estimates correspond to instantaneous decay rates per generation of 0.019 (= 0.0045 × 4.2) for age-4 egg size and 0.024 for age-5 egg size. In contrast, the corresponding instantaneous per generation decay rates were approximately 0.26 for return rate and 0.36 for chum salmon abundance after the breakpoint. A previous study based on steelhead (*O. mykiss*), brown trout (*S. trutta*), and farmed Atlantic salmon predicted declines in relative reproductive success (RRS) with an exponential instantaneous decay rate of *b̂* = 0.375 per generation (Araki et al. 2007). The estimated instantaneous decay rate for chum salmon abundance after the breakpoint (0.36 per generation) was remarkably close to the RRS-based estimate of 0.375 per generation from Araki et al. (2007), corresponding to declines of approximately 30% and 31% per generation, respectively. The estimated rate for return rate per generation (0.26) was lower but still broadly consistent with the RRS-based prediction.

Mean female fork length (FL) was positively associated with mean egg size across brood years (*r* = 0.80), indicating that larger females generally had larger egg sizes and explaining 64% of the variation in mean egg size (Fig. 7A). Mean egg size of age-4 and age-5 females was positively correlated with brood-year-specific return rates of hatchery-reared chum salmon (ages 2–5) (*r* = 0.70), explaining 46% of the variation in return rate (Fig. 7B).

**Fig. 6.**
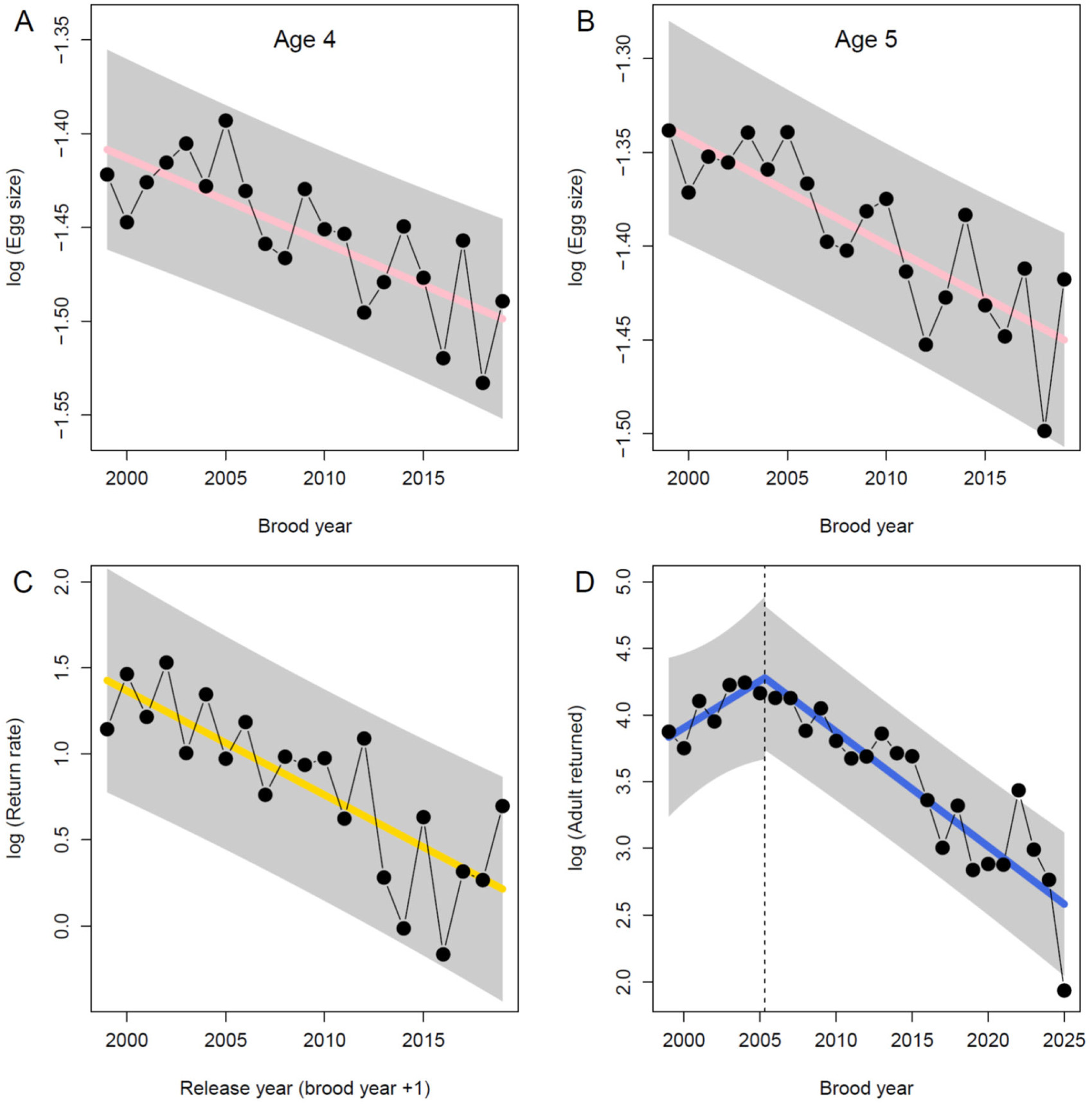
Temporal trends in egg size, return rate, and abundance of Japanese chum salmon. Exponential decay models were fitted as *N* = *ae*^−*bx*^ (see Table 1). (A) Egg size of age-4 fish, (B) age-5 fish, (C) return rate, ages-2–5 (1999–2019), and (D) Number of chum salmon returning to Japan (1999-2025). Filled circles indicate observed values, solid lines show fitted values, and shaded areas represent 95% prediction intervals. Panels A–C show exponential decay models, whereas panel D shows a segmented regression with an estimated breakpoint in 2005.3, marked by the dashed vertical line (Table 1).

**Fig. 7.**
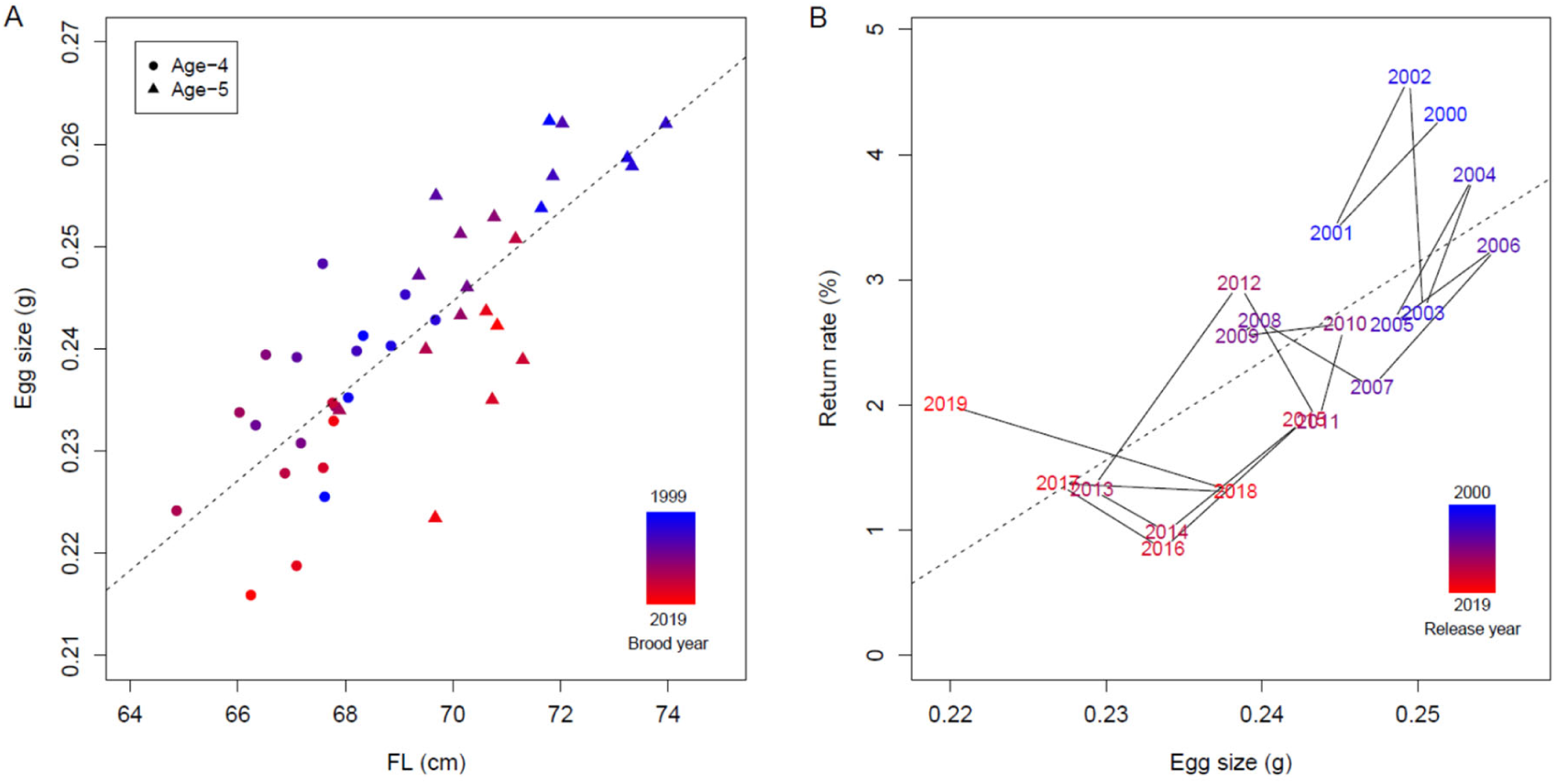
Relationships between egg size, return rate, and female fork length (FL). (A) Mean FL versus mean egg size for ages-4, 5 for each brood year. The dashed line indicates the linear regression based on combined data of ages-4, 5 (*r* = 0.8, adjusted *R*² = 0.64, *p* < 0.001). (B) Mean egg size ages 4 and 5 versus return rate (ages-2–5) of hatchery-reared chum salmon originating from each brood year. Each point is labeled with the release year (cohort) (brood year + 1). The dashed line indicates the linear regression (*r* = 0.7, adjusted *R*² = 0.46, *p* < 0.001).

## 4. Discussion

The analyses revealed a widespread long-term decline in egg size across 13 hatchery-enhanced chum salmon populations in Japan from 1999 to 2019. Despite substantial within- and among-river variation, the decline occurred synchronously across geographically separated populations, suggesting that broad-scale environmental or biological processes have influenced reproductive traits. In contrast, female FL showed pronounced interannual fluctuations but no consistent nationwide decline, and the persistent decline in egg size after accounting for female FL, age, and river effects indicates that reduced egg size cannot be explained solely by changes in maternal body size. Nevertheless, female FL remained strongly and positively correlated with egg size (*r* = 0.80), highlighting the importance of body size in reproductive investment. Nationwide mean egg size was positively associated with subsequent return rate (*r* = 0.70), suggesting that long-term changes in reproductive investment may be linked to declining post-release survival and population performance. The concurrent shift toward younger ages at return further indicates that multiple fitness-related life-history traits have changed during the period of population decline.

### 4.1 Temporal variation in female body size

The global smooth for female FL from the river-specific GAM showed a temporal pattern broadly consistent with that of the hierarchical GAM, despite substantial among-river and individual variation. The hierarchical GAM, which accounted for age and river effects, revealed pronounced interannual fluctuations in female FL (Fig. 4A), with peaks around 2003, 2009, and 2015. After reaching a minimum around 2012, female fork length increased and subsequently fluctuated around a relatively stable level. Similar temporal variability in salmon body size has been linked to climate-driven changes in marine growth conditions and increasing competition at sea in previous studies (Lewis et al. 2015; Jeffrey et al. 2017; Oke et al. 2020; Ohlberger et al. 2023). Although the mechanisms underlying the observed fluctuations in female FL of Japanese chum salmon remain unresolved, they are likely to reflect complex interactions among environmental variation, density-dependent processes, and changes in the North Pacific marine ecosystem.

### 4.2 Long-term decline in egg size after accounting for female body size

The global smooth from the river-specific GAM showed a decline in egg size that was broadly consistent across river populations. The hierarchical GAM demonstrated a persistent decline in egg size across rivers even after accounting for the effects of female FL, age, and river (Fig. 4B). The decline remained evident despite substantial interannual fluctuations in female FL, indicating that factors beyond temporal variation in female body size contributed to the long-term reduction in egg size.

Female body size is generally regarded as one of the strongest determinants of egg size in salmonids, with larger females typically producing larger eggs (Beacham and Murray 1985, 1993; Fleming and Gross 1990; Einum and Fleming 1999, 2000; Heath et al. 1999). Consistent with this general pattern, nationwide annual mean FL was strongly correlated with mean egg size (*r* = 0.80) in the present study (Fig. 7A). Nevertheless, the persistence of the declining trend after statistically controlling for FL indicates that factors other than maternal size have contributed to the long-term reduction in egg size. This finding suggests that changes in reproductive allocation, rather than changes in female body size alone, may underlie the observed decline in egg size.

### 4.3 Shift toward younger age at maturity

The age composition analysis revealed that the proportion of age-6 and age-5 fish declined, whereas the proportions of age-3 and age-4 fish increased, albeit with considerable interannual variation, resulting in a marked decline in mean age at return after approximately 2010 (Fig. 5).

Shifts toward younger age at maturity have been reported in Pacific salmon populations, where declines in body size and age at maturity have been linked to climate-driven changes in marine conditions (Lewis et al. 2015; Oke et al. 2020; Ohlberger et al. 2023). Comparable patterns have also been documented in Atlantic salmon populations, where long-term declines in age and body size at maturity have been associated with environmental change (Besnier et al. 2024; Fiske et al. 2025). Besnier et al. (2024) further showed that recent environmental changes have weakened the influence of major maturation genes (*vgll3* and *six6*) through growth-driven plasticity. In hatchery-reared steelhead, Harris et al. (2026) reported changes in adaptive genetic variation near the *six6* locus, which is associated with age at maturity and body size, without a corresponding loss of neutral genetic diversity. Similar patterns have been reported for Japanese chum salmon also retain high levels of neutral genetic diversity across the Pacific Rim (Beacham et al. 2008; Kitada 2018), whereas allele frequencies at many adaptive SNP loci differ from expectations based on neutral population structure, suggesting that hatchery-related selection may have contributed to these patterns (Kitada and Kishino 2021). Furthermore, hatchery programs in Hokkaido have increasingly focused on early-run populations since the early 1980s, and late-run populations had largely disappeared by the late 1990s (Miyakoshi et al. 2013). Whether these genetic changes and hatchery management practices have contributed to the observed shift toward younger age at maturity in Japanese chum salmon remains unknown.

The timing of the shift toward younger age at return differed from that of the decline in abundance. Segmented regression indicated a marked change in the rate of abundance decline around 2005 (Fig. 6D), whereas the decline in mean age at return became apparent only after approximately 2010 (Fig. 5). This temporal difference may indicate that reduced post-release survival or other environmental effects on abundance preceded detectable changes in age at maturity. Alternatively, the lag may reflect the time required for changes in growth, maturation, or their underlying selective pressures to become evident in the age composition. Although the mechanisms cannot be distinguished from the present data, the temporal sequence suggests that the processes underlying declining abundance and younger age at return may not have operated synchronously.

### 4.4 Egg-size decline and lifetime fitness

Hatchery propagation can relax natural selection favoring large eggs and promote rapid evolutionary shifts toward smaller egg size (Heath et al. 2003). In Japanese chum salmon, however, egg size declined much more slowly, by only approximately 2% per generation. The average return rate of Japanese hatchery chum salmon was 2.5 ± 1.1% for the 1999– 2019 brood years (Kitada et al. 2025), suggesting that strong natural selection in the wild after release remains a dominant evolutionary force. Such strong post-release selection may partially counteract the relaxation of selection during hatchery incubation and rearing, where smaller eggs may survive that would otherwise be removed by natural selection, thereby contributing to the relatively gradual decline in egg size compared with the marked declines in abundance and return rate.

Despite the much smaller absolute decline in egg size, the instantaneous decay rates estimated from standardized variables were remarkably similar among egg size, return rate, and chum salmon abundance (Table 1). Notably, egg size peaked in 2005 and declined thereafter (Fig. 4B), coinciding with the estimated breakpoint (2005.3) in chum salmon abundance, after which abundance shifted from an increasing to a declining trend (Fig. 6D). These concordant rates of decline and the temporal coincidence of the egg size peak and abundance breakpoint, together with the positive relationship between egg size and return rate (Fig. 7B), suggest that long-term declines in egg size may be closely associated with population decline in Japanese chum salmon. Because egg size is a central life-history trait reflecting the trade-off between fecundity and offspring size, evolutionary changes in this trait may have consequences that extend beyond early survival, potentially influencing lifetime fitness and population dynamics.

### 4.5 Evolutionary implications of long-term hatchery propagation

My findings of long-term changes in female body size, egg size, and age at return are consistent with growing evidence that large-scale artificial propagation can alter life-history traits and fitness-related characteristics in Pacific salmon (Reisenbichler and Rubin 1999; Heath et al. 2003; Araki et al. 2007; Christie et al. 2012a; Christie et al. 2014; Shedd et al. 2022; Malick et al. 2023) and Atlantic salmon (Fleming et al. 2000; McGinnity et al. 2003; Fiske et al. 2025). However, the mechanisms underlying these changes are likely to differ between species because of differences in both life history and propagation practices. Pacific salmon are predominantly semelparous, whereas Atlantic salmon are iteroparous. In Atlantic salmon, farmed fish have undergone substantial domestication selection associated with aquaculture, particularly for growth-related traits (Næve et al. 2022), and escaped juveniles and adults can reproduce with wild fish, making genetic introgression a major conservation concern (Taranger et al. 2015; Glover et al. 2017; Bolstad et al. 2021; Diserud et al. 2022; San Román et al. 2025). In contrast, Japanese chum salmon have been maintained by large-scale hatchery supplementation for more than 130 years (Fig. S1). Although natural spawning by chum salmon has been documented in Japanese rivers, most returning chum salmon are hatchery-released fish or wild-born hatchery descendants (Kitada 2020). Consequently, genetic introgression from hatchery fish into wild populations is likely to play a much smaller role than in Atlantic salmon. Instead, the long-term changes observed here are more likely to reflect evolutionary and demographic changes within the hatchery population itself.

The estimated instantaneous exponential decay rate for chum salmon abundance after the breakpoint (0.36 per generation) was close to the RRS-based estimate of 0.375 per generation from Araki et al. (2007), although these quantities are not directly equivalent. The abundance-based estimate reflects the decline in post-release survival from release to return of hatchery fish returning in a given year (across ages), whereas the RRS-based estimate reflects the decline in lifetime fitness from egg to return of hatchery-origin fish born in a given year in the wild. RRS-based estimates are influenced by variation in spawning river fidelity, reproductive success rate and survival during the early life stages, including the critical period from egg to fry. In hatchery-reared chum salmon, survival through this critical period occurs largely under hatchery conditions and is therefore very high, whereas mortality of fry after release in coastal waters can be substantial (Kitada et al. 2025). Notably, the temporal scale of decline also differs among species and measures of fitness. In steelhead, declines in RRS have been detected from a single generation of hatchery rearing (Araki et al. 2007; Christie et al. 2012a), whereas chum salmon abundance began to decline approximately three decades, corresponding to about seven generations, after the onset of large-scale hatchery propagation in the mid-1970s (Fig. S1). This contrast highlights that changes in reproductive success and population abundance may emerge on different temporal scales under prolonged hatchery propagation. Thus, the similarity between the two estimates should be interpreted cautiously, as they capture different components and temporal scales of population performance.

Return rate also measures the post-release survival of hatchery fish from release to return across ages for the corresponding release cohort. The estimated instantaneous decay rate for chum salmon return rate was 0.26 per generation. In contrast, the corresponding rate for hatchery-propagated red sea bream (*Pagrus major*) was much higher, at 0.14 per year, equivalent to 0.67 per generation based on a mean generation time of 4.8 years (Kitada et al. 2019). Red sea bream were propagated from a relatively small captive broodstock of approximately 130 adults, although wild fish were occasionally incorporated into the broodstock. Catches of hatchery-released fish increased rapidly, reaching a maximum in 1991, 17 years, corresponding to approximately 3.5 generations, after the program began. In contrast, Japanese chum salmon have been propagated primarily from large numbers of returning hatchery adults that had undergone natural selection after release, and chum salmon abundance peaked in 2004, approximately seven generations after large-scale hatchery release. Thus, in both species, the initial increase in returns following the expansion of hatchery releases was followed by a long-term decline in hatchery fish despite continued hatchery propagation. This common temporal pattern, although based on only two examples, raises the possibility that prolonged hatchery propagation can eventually be followed by substantial declines in population-level fitness, with the magnitude and timing of such changes potentially influenced by broodstock size and source, the number of generations of artificial propagation, and environmental variation. Species-specific life-history differences may also contribute: red sea bream produce approximately 1.9 million small eggs (mean diameter 0.9 mm), whereas chum salmon produce only about 2,600–3,200 much larger eggs (7.9–8.6 mm in diameter) (Matsuura et al. 1988; Beacham and Murray 1993).

Finally, the observed return in 2025 fell below the prediction interval of the fitted model (Fig. 6D), suggesting that recent marine conditions may have become more unfavorable than those represented by the historical trend. Whether this deviation reflects an unusually poor return year or the onset of a more persistent decline remains uncertain. However, the long-term decline observed in other hatchery-enhanced species, such as red sea bream, highlights the importance of continued monitoring to determine whether recent changes represent short-term environmental fluctuations or more persistent changes in fitness-related traits and population performance.

## 5. Conclusions

This study revealed widespread long-term changes in key life-history traits across hatchery-enhanced chum salmon populations in Japan. Hierarchical GAM analyses demonstrated a nationwide decline in egg size after accounting for variation in female body size, age, and river effects, indicating that the decline cannot be explained solely by changes in maternal size. In addition, mean age at return declined following shifts in age composition, providing further evidence of long-term changes in the life-history schedule of Japanese chum salmon. Egg size was positively associated with return rate, and standardized exponential decay rates were remarkably similar among egg size, return rate, and population abundance, suggesting that these long-term declines may be driven by interacting environmental and evolutionary processes.

Together, these findings demonstrate that fitness-related life-history traits of Japanese chum salmon have changed substantially over the past two decades and suggest that these changes may be associated with declining lifetime fitness and population abundance. Although the relative contributions of environmental change, hatchery selection, and evolutionary responses remain unresolved, integrating long-term phenotypic, environmental, and genomic data will be essential for identifying the mechanisms underlying these changes.

Beyond Japanese chum salmon, this study highlights the need to evaluate hatchery propagation not only by short-term demographic outcomes but also by its potential long-term evolutionary consequences. Understanding how artificial propagation influences life-history traits and lifetime fitness will be essential for developing sustainable enhancement and conservation strategies that maintain fish abundance while preserving evolutionary resilience and minimizing unintended evolutionary consequences.

## Author contributions

Shuichi Kitada; Conceptualization, Data curation, Formal analysis, Funding acquisition, Methodology, Software, Validation, Visualization, Writing – original draft, Writing – review and editing.

## Acknowledgements

I am grateful to Hirohisa Kishino for valuable suggestions and in-depth discussions during the early stages of this study, which helped shape the manuscript. I sincerely thank Trevor Pitcher, Editor-in-Chief, for overseeing the editorial process, and the anonymous Associate Editor and two anonymous reviewers for their insightful and constructive comments throughout the review process. I thank the many biologists, fisheries and hatchery managers, and fishers who contributed to the collection of the data analyzed in this study. I also acknowledge the efforts of the Fisheries Research and Education Agency (FRA) and the North Pacific Anadromous Fish Commission (NPAFC) in developing and maintaining the publicly available databases that made this research possible. ChatGPT (OpenAI, GPT-5.5) was used to assist with English editing, manuscript preparation, and code troubleshooting. All scientific interpretations, analyses, and conclusions were developed and verified by the author. This work was supported by the Japan Society for the Promotion of Science (JSPS) through Grants-in-Aid for Scientific Research (KAKENHI) (No. 18K05781).

## Conflict of interest

The authors declare that no conflict of interest exists.

## Data availability statement

The data generated or analyzed in this study are fully described in the published article and its Supplementary Information. Analysis scripts and data will be available on acceptance.

## Notes

### Competing Interest Statement

The authors have declared no competing interest.

### Summary of Updates

1.Additional segmented regression analyses. Figure 6D and Table 1 were updated. 2.Individual observations were removed from the river-specific GAM figures, and global smooths were added (Figures 2, 3, S6, and S7). Residual diagnostic checks were also conducted, with results provided in new Supplementary Figures S8-S11. 3.Figure 5 was revised to include a GAM-smoothed line for the annual weighted mean age at return. 4.Substantial revision of Discussion Section 4.5.

