## Supplementary Information for "Long-term decline in egg size of Japanese hatchery chum salmon links to declines in return rate and abundance"

Shuichi Kitada[1*]

[1] Tokyo University of Marine Science and Technology, Tokyo, Japan

* Kitada(at)kaiyodai.ac.jp

**This PDF file includes:**
Tables S1–S16
Figures S1–S17

**Table S1. Sample data of hatchery-reared chum salmon collected from 13 rivers in Japan, including measurements of female.**

| | River | Latitude N (river mouth) | Sampling year | Age-4 | | | Age-5 | | |
|---|---|---|---|---|---|---|---|---|---|
| | | | | n | Female FL ± SD (cm) | Egg size ± SD (g) | n | Female FL ± SD (cm) | Egg size ± SD (g) |
| 1 | Tokushibetsu | 44°50' | 1997–2020 | 1,108 | 65.85 ± 3.47 | 0.237 ± 0.030 | 817 | 69.32 ± 3.74 | 0.251 ± 0.032 |
| 2 | Ishikari | 43°15' | 1997–2020 | 1,307 | 65.69 ± 3.39 | 0.199 ± 0.025 | 613 | 68.50 ± 3.58 | 0.209 ± 0.030 |
| 3 | Nishibetsu | 43°22' | 1997–2020 | 1,303 | 64.70 ± 3.46 | 0.240 ± 0.031 | 794 | 67.62 ± 3.85 | 0.253 ± 0.038 |
| 4 | Tokachi | 42°41' | 1997–2020 | 1,152 | 66.34 ± 3.81 | 0.268 ± 0.034 | 1,131 | 70.04 ± 3.96 | 0.289 ± 0.039 |
| 5 | Yurappu | 42°16' | 1997–2020 | 1,060 | 69.17 ± 3.34 | 0.257 ± 0.030 | 1,072 | 72.88 ± 3.72 | 0.274 ± 0.035 |
| 6 | Akka | 40° 3' | 1999–2020 | 877 | 65.95 ± 3.69 | 0.241 ± 0.032 | 674 | 70.16 ± 4.17 | 0.260 ± 0.036 |
| 7 | Tsugaruishi | 39°35' | 1997–2019 | 922 | 70.53 ± 3.80 | 0.251 ± 0.033 | 1,062 | 75.02 ± 4.17 | 0.273 ± 0.039 |
| 8 | Katagishi | 39°12' | 1997–2019 | 1,048 | 66.49 ± 3.87 | 0.238 ± 0.031 | 839 | 70.34 ± 4.24 | 0.251 ± 0.035 |
| 9 | Kitakami | 39°34' | 1997–2020 | 1,160 | 66.57 ± 3.62 | 0.229 ± 0.032 | 589 | 69.81 ± 3.77 | 0.239 ± 0.035 |
| 10 | Uda | 37°48' | 1999–2020 | 894 | 66.76 ± 3.46 | 0.216 ± 0.031 | 468 | 69.30 ± 3.95 | 0.231 ± 0.032 |
| 11 | Gakko | 39° 4' | 1997–2020 | 1,087 | 70.53 ± 3.81 | 0.233 ± 0.032 | 707 | 74.67 ± 3.93 | 0.250 ± 0.035 |
| 12 | Miomote | 38°14' | 1999–2019 | 1,043 | 69.75 ± 3.59 | 0.232 ± 0.029 | 451 | 72.70 ± 3.89 | 0.243 ± 0.034 |
| 13 | Sho | 36°47' | 1997–2020 | 1,123 | 68.39 ± 3.70 | 0.220 ± 0.028 | 366 | 71.83 ± 4.01 | 0.232 ± 0.032 |

**Table S2.** Sample sizes of age-4 female chum salmon by river and year for 13 rivers in Japan. NA: not available. See Fig. 1 for river locations.

| Brood year | Tokushi-betsu | Ishikari | Nishi-betsu | Tokachi | Yurappu | Akka | Tsugaru-ishi | Kata-gishi | Kita-kami | Uda | Gakko | Mio-mote | Sho | Total |
|---|---|---|---|---|---|---|---|---|---|---|---|---|---|---|
| 1997 | 111 | 116 | 125 | 82 | 48 | NA | 59 | 25 | 123 | NA | 23 | NA | 77 | 789 |
| 1998 | 72 | 76 | 73 | 81 | 68 | NA | 28 | 54 | 73 | NA | NA | NA | 26 | 551 |
| 1999 | 27 | 52 | 50 | 15 | 29 | 6 | 24 | 5 | 40 | 45 | 65 | 42 | 74 | 474 |
| 2000 | 75 | 82 | 66 | 76 | 88 | 84 | 37 | 64 | 57 | NA | 125 | 39 | 43 | 836 |
| 2001 | 31 | 38 | 66 | 37 | 97 | 45 | 65 | 58 | 55 | 54 | 33 | 21 | 20 | 620 |
| 2002 | 61 | 80 | 65 | 146 | 28 | 12 | 48 | 34 | NA | 72 | 51 | 58 | 88 | 743 |
| 2003 | NA | 50 | 62 | 47 | 78 | 69 | 18 | 69 | 36 | 38 | 42 | 44 | 41 | 594 |
| 2004 | 63 | 85 | 21 | 36 | 17 | 50 | 34 | 74 | 55 | 25 | 90 | 60 | NA | 610 |
| 2005 | 63 | 35 | 88 | 62 | 57 | 69 | 62 | 48 | 75 | 79 | 20 | 62 | 67 | 787 |
| 2006 | 35 | 59 | 26 | 9 | 40 | 21 | 25 | 54 | 37 | 58 | 40 | 58 | 39 | 501 |
| 2007 | 21 | 40 | 81 | 83 | 47 | 71 | 63 | 65 | 40 | 76 | 37 | 68 | 79 | 771 |
| 2008 | 47 | 31 | 49 | 45 | 16 | 44 | 28 | 47 | 49 | 53 | 62 | 54 | 44 | 569 |
| 2009 | 82 | 57 | 55 | 43 | 66 | 50 | 56 | 38 | 64 | 19 | 59 | 83 | 72 | 744 |
| 2010 | 41 | 50 | 60 | 18 | 42 | 48 | 45 | 30 | 45 | 63 | 29 | 51 | 35 | 557 |
| 2011 | 40 | 81 | 59 | 40 | 51 | 61 | 23 | 53 | 60 | NA | 44 | 65 | 55 | 632 |
| 2012 | 44 | 64 | 49 | 34 | 38 | 53 | 37 | 50 | 43 | 32 | 64 | 50 | 46 | 604 |
| 2013 | 57 | 20 | 51 | 74 | 8 | 49 | 43 | 36 | 68 | 48 | 42 | 49 | 65 | 610 |
| 2014 | 65 | 49 | 41 | 37 | 49 | 12 | 3 | 14 | 52 | 16 | 53 | 67 | 44 | 502 |
| 2015 | 46 | 70 | 74 | 52 | 65 | 64 | 42 | 37 | 31 | 86 | 60 | 68 | 70 | 765 |
| 2016 | 18 | 37 | 12 | 24 | 5 | NA | 57 | 44 | 32 | 30 | 26 | 19 | 33 | 337 |
| 2017 | 23 | 33 | 30 | 5 | 30 | 21 | 45 | 54 | 31 | 24 | 32 | 42 | 36 | 406 |
| 2018 | 22 | 29 | 33 | 43 | 30 | 40 | 66 | 80 | 40 | 33 | 39 | 23 | 38 | 516 |
| 2019 | 31 | 31 | 33 | 32 | 27 | 2 | 14 | 15 | 13 | NA | 14 | 20 | 26 | 258 |
| 2020 | 33 | 42 | 34 | 31 | 36 | 6 | NA | NA | 41 | 43 | 37 | NA | 5 | 308 |
| Total | 1,108 | 1,307 | 1,303 | 1,152 | 1,060 | 877 | 922 | 1,048 | 1,160 | 894 | 1,087 | 1,043 | 1,123 | 14,084 |

**Table S3.** Sample sizes of age-5 female chum salmon by river and year for 13 rivers in Japan. NA: not available. See Fig. 1 for river locations.

| Brood year | Tokushi-betsu | Ishikari | Nishi-betsu | Tokachi | Yurappu | Akka | Tsugaru-ishi | Kata-gishi | Kita-kami | Uda | Gakko | Mio-mote | Sho | Total |
|---|---|---|---|---|---|---|---|---|---|---|---|---|---|---|
| 1997 | 51 | 34 | 33 | 88 | 114 | NA | 24 | 27 | 27 | NA | 11 | NA | 12 | 421 |
| 1998 | 27 | 14 | 17 | 17 | 25 | NA | 37 | 29 | 42 | NA | NA | NA | 2 | 210 |
| 1999 | 69 | 39 | 50 | 78 | 60 | 41 | 10 | 34 | 28 | 29 | 60 | 6 | 10 | 514 |
| 2000 | 19 | 11 | 30 | 9 | 4 | 4 | 42 | 7 | 17 | NA | 17 | 28 | 11 | 199 |
| 2001 | 64 | 41 | 30 | 61 | 76 | 49 | 21 | 18 | 21 | 4 | 25 | 12 | 33 | 455 |
| 2002 | 35 | 17 | 29 | 145 | 62 | 72 | 36 | 45 | NA | 10 | 37 | 24 | 5 | 517 |
| 2003 | NA | 31 | 33 | 51 | 14 | 16 | 75 | 13 | 10 | 24 | 30 | 13 | 41 | 351 |
| 2004 | 34 | 9 | 75 | 60 | 78 | 43 | 60 | 60 | 20 | 45 | 8 | 11 | NA | 503 |
| 2005 | NA | 57 | 7 | 33 | 30 | 25 | 28 | 44 | 15 | 7 | 76 | 22 | 18 | 362 |
| 2006 | 61 | 33 | 64 | 83 | 57 | 41 | 69 | 39 | 51 | 31 | 42 | 30 | 34 | 635 |
| 2007 | 62 | 52 | 11 | 14 | 47 | 16 | 29 | 27 | 20 | 20 | 15 | 26 | 12 | 351 |
| 2008 | 34 | 42 | 48 | 51 | 70 | 48 | 66 | 46 | 29 | 33 | 28 | 28 | 24 | 547 |
| 2009 | 16 | 38 | 40 | 54 | 30 | 41 | 40 | 46 | 24 | 56 | 36 | 5 | 11 | 437 |
| 2010 | 57 | 32 | 28 | 67 | 51 | 42 | 54 | 63 | 38 | 7 | 69 | 34 | 58 | 600 |
| 2011 | 53 | 3 | 40 | 49 | 41 | 33 | 54 | 31 | 30 | NA | 40 | 27 | 22 | 423 |
| 2012 | 49 | 32 | 50 | 62 | 59 | 40 | 58 | 38 | 44 | 54 | 23 | 31 | 6 | 546 |
| 2013 | 31 | 71 | 45 | 23 | 83 | 37 | 53 | 51 | 18 | 42 | 54 | 51 | 8 | 567 |
| 2014 | 28 | 18 | 46 | 59 | 27 | 59 | 92 | 58 | 44 | 61 | 28 | 26 | 13 | 559 |
| 2015 | 47 | 4 | 23 | 39 | 32 | 4 | 31 | 28 | 50 | 6 | 35 | 27 | 10 | 336 |
| 2016 | 21 | 11 | 36 | 19 | 43 | NA | 28 | 42 | 13 | 17 | 21 | 14 | 9 | 274 |
| 2017 | 18 | 8 | 15 | 40 | 15 | 4 | 49 | 33 | 5 | 9 | 14 | 2 | 7 | 219 |
| 2018 | 24 | 12 | 13 | 4 | 20 | 10 | 33 | 11 | 8 | 10 | 10 | 25 | 8 | 188 |
| 2019 | 10 | 1 | 17 | 11 | 22 | 48 | 73 | 49 | 32 | NA | 26 | 9 | 11 | 309 |
| 2020 | 7 | 3 | 14 | 14 | 12 | 1 | NA | NA | 3 | 3 | 2 | NA | 1 | 60 |
| Total | 817 | 613 | 794 | 1,131 | 1,072 | 674 | 1,062 | 839 | 589 | 468 | 707 | 451 | 366 | 9,583 |

**Table S4.** Parametric coefficients of the river-specific generalized additive model (GAM) for age-4 female fork length (FL) after accounting for river effects; *FL* ~ *s*(year) + *s*(year, by = river) + river (Fig. 2). River coefficients represent differences relative to the reference river (Tokushibetsu). The river-specific GAM for age-4 female FL explained 31.3% of the deviance (adjusted $R^2$ = 0.31).

| Coefficient | Estimate | Standard error | *t* | *p*-value |
|---|---|---|---|---|
| (Intercept) | 65.8814 | 0.1136 | 580.01 | **< 0.0001** |
| riverIshikari | -0.2682 | 0.1534 | -1.75 | 0.0805 |
| riverNishibetsu | -1.0100 | 0.1538 | -6.57 | **< 0.0001** |
| riverTokachi | 0.4053 | 0.1583 | 2.56 | **0.0105** |
| riverYurappu | 3.2492 | 0.1604 | 20.26 | **< 0.0001** |
| riverAkka | 0.1557 | 0.1628 | 0.96 | 0.3390 |
| riverTsugaruishi | 4.7533 | 0.1660 | 28.63 | **< 0.0001** |
| riverKatagishi | 0.7053 | 0.1591 | 4.43 | **< 0.0001** |
| riverKitakami | 0.6038 | 0.1602 | 3.77 | **0.0002** |
| riverUda | 0.7817 | 0.1630 | 4.80 | **< 0.0001** |
| riverGakko | 4.7869 | 0.1565 | 30.58 | **< 0.0001** |
| riverMiomote | 3.9969 | 0.1559 | 25.64 | **< 0.0001** |
| riverSho | 2.4936 | 0.1563 | 15.96 | **< 0.0001** |

**Table S5.** Approximate significance of smooth terms in the river-specific generalized additive model (GAM) for age-4 female fork length (FL) after accounting for river effects; *FL* ~ *s*(year) + *s*(year, by = river) + river (Fig. 2). Non-significant river-specific smooth terms indicate that temporal trends in those rivers did not differ significantly from the overall smooth trend.

| Smooth term | edf | Ref. df | *F* | *p*-value |
|---|---|---|---|---|
| s(year) | 8.791 | 8.964 | 40.77 | **< 0.0001** |
| s(year):riverTokushibetsu | 2.534 | 3.156 | 2.52 | 0.0506 |
| s(year):riverIshikari | 1.045 | 1.086 | 3.49 | 0.0543 |
| s(year):riverNishibetsu | 8.576 | 8.944 | 11.32 | **< 0.0001** |
| s(year):riverTokachi | 1.003 | 1.006 | 1.59 | 0.2077 |
| s(year):riverYurappu | 2.801 | 3.484 | 2.18 | 0.0629 |
| s(year):riverAkka | 2.990 | 3.884 | 3.59 | **0.0047** |
| s(year):riverTsugaruishi | 7.192 | 8.202 | 5.14 | **< 0.0001** |
| s(year):riverKatagishi | 2.689 | 3.334 | 3.07 | **0.0224** |
| s(year):riverKitakami | 5.506 | 6.650 | 2.84 | **0.0105** |
| s(year):riverUda | 1.028 | 1.055 | 4.44 | **0.0346** |
| s(year):riverGakko | 7.065 | 8.110 | 6.77 | **< 0.0001** |
| s(year):riverMiomote | 5.210 | 6.311 | 2.57 | **0.0152** |
| s(year):riverSho | 5.181 | 6.257 | 2.79 | **0.0110** |

edf, estimated degrees of freedom; Ref. df, reference degrees of freedom used in the approximate *F* test for smooth terms.

**Table S6.** Parametric coefficients of the river-specific generalized additive model (GAM) for age-4 egg size after accounting for river effects; *Egg* ~ *s*(year) + *s*(year, by = river) + river (Fig. 3). River coefficients represent differences relative to the reference river (Tokushibetsu). The river-specific GAM for age-4 egg size explained 32.1% of the deviance (adjusted $R^2 = 0.32$).

| Coefficient | Estimate | Standard error | *t* | *p*-value |
|---|---|---|---|---|
| (Intercept) | 0.2333 | 0.0010 | 238.93 | < **0.0001** |
| riverIshikari | -0.0359 | 0.0013 | -27.29 | < **0.0001** |
| riverNishibetsu | 0.0063 | 0.0013 | 4.82 | < **0.0001** |
| riverTokachi | 0.0331 | 0.0014 | 23.88 | < **0.0001** |
| riverYurappu | 0.0219 | 0.0014 | 16.01 | < **0.0001** |
| riverAkka | 0.0073 | 0.0014 | 5.25 | < **0.0001** |
| riverTsugaruishi | 0.0187 | 0.0014 | 13.33 | < **0.0001** |
| riverKatagishi | 0.0053 | 0.0013 | 3.91 | < **0.0001** |
| riverKitakami | -0.0062 | 0.0014 | -4.53 | < **0.0001** |
| riverUda | -0.0179 | 0.0014 | -12.88 | < **0.0001** |
| riverGakko | 0.0011 | 0.0013 | 0.83 | 0.405 |
| riverMiomote | -0.0009 | 0.0013 | -0.66 | 0.507 |
| riverSho | -0.0143 | 0.0013 | -10.72 | < **0.0001** |

**Table S7.** Approximate significance of smooth terms in the river-specific generalized additive model (GAM) for age-4 egg size after accounting for river effects; *Egg* ~ *s*(year) + *s*(year, by = river) + river (Fig. 3). Non-significant river-specific smooth terms indicate that temporal trends in those rivers did not differ significantly from the overall smooth trend.

| Smooth term | edf | Ref. df | *F* | *p*-value |
|---|---|---|---|---|
| s(year) | 7.863 | 8.566 | 10.76 | < **0.0001** |
| s(year):riverTokushibetsu | 7.805 | 8.607 | 9.11 | < **0.0001** |
| s(year):riverIshikari | 4.934 | 5.995 | 1.43 | 0.1865 |
| s(year):riverNishibetsu | 7.543 | 8.452 | 5.31 | < **0.0001** |
| s(year):riverTokachi | 8.670 | 8.961 | 9.82 | < **0.0001** |
| s(year):riverYurappu | 1.014 | 1.028 | 0.69 | 0.4008 |
| s(year):riverAkka | 4.414 | 5.372 | 2.57 | **0.0297** |
| s(year):riverTsugaruishi | 2.405 | 2.986 | 2.96 | **0.0332** |
| s(year):riverKatagishi | 1.006 | 1.012 | 1.39 | 0.2374 |
| s(year):riverKitakami | 8.068 | 8.750 | 6.09 | < **0.0001** |
| s(year):riverUda | 0.006 | 0.011 | 0.07 | 0.9779 |
| s(year):riverGakko | 7.746 | 8.575 | 6.67 | < **0.0001** |
| s(year):riverMiomote | 1.014 | 1.027 | 0.22 | 0.652 |
| s(year):riverSho | 5.466 | 6.558 | 7.67 | < **0.0001** |

edf, estimated degrees of freedom; Ref. df, reference degrees of freedom used in the approximate *F* test for smooth terms.

**Table S8.** Parametric coefficients of the river-specific generalized additive model (GAM) for age-5 female fork length (FL) after accounting for river effects; *FL* ~ *s*(year) + *s*(year, by = river) + river (Fig. S6). River coefficients represent differences relative to the reference river (Tokushibetsu). The river-specific GAM for age-5 female FL explained 38.0% of the deviance (adjusted $R^2$ = 0.38).

| Coefficient | Estimate | Standard error | *t* | *p*-value |
|---|---|---|---|---|
| (Intercept) | 0.2333 | 0.0010 | 238.93 | < **0.0001** |
| riverIshikari | -0.0359 | 0.0013 | -27.29 | < **0.0001** |
| riverNishibetsu | 0.0063 | 0.0013 | 4.82 | < **0.0001** |
| riverTokachi | 0.0331 | 0.0014 | 23.88 | < **0.0001** |
| riverYurappu | 0.0219 | 0.0014 | 16.01 | < **0.0001** |
| riverAkka | 0.0073 | 0.0014 | 5.25 | < **0.0001** |
| riverTsugaruishi | 0.0187 | 0.0014 | 13.33 | < **0.0001** |
| riverKatagishi | 0.0053 | 0.0013 | 3.91 | < **0.0001** |
| riverKitakami | -0.0062 | 0.0014 | -4.53 | < **0.0001** |
| riverUda | -0.0179 | 0.0014 | -12.88 | < **0.0001** |
| riverGakko | 0.0011 | 0.0013 | 0.83 | 0.405 |
| riverMiomote | -0.0009 | 0.0013 | -0.66 | 0.507 |
| riverSho | -0.0143 | 0.0013 | -10.72 | < **0.0001** |

**Table S9.** Approximate significance of smooth terms in the river-specific generalized additive model (GAM) for age-5 female fork length (FL) after accounting for river effects; *FL* ~ *s*(year) + *s*(year, by = river) + river (Fig. S6). Non-significant river-specific smooth terms indicate that temporal trends in those rivers did not differ significantly from the overall smooth trend.

| Smooth term | edf | Ref. df | *F* | *p*-value |
|---|---|---|---|---|
| s(year) | 8.856 | 8.978 | 51.93 | **< 0.0001** |
| s(year):riverTokushibetsu | 2.680 | 3.331 | 3.07 | **0.0206** |
| s(year):riverIshikari | 5.225 | 6.287 | 3.29 | **0.0030** |
| s(year):riverNishibetsu | 6.982 | 8.024 | 4.50 | **< 0.0001** |
| s(year):riverTokachi | 7.732 | 8.568 | 3.02 | **0.0268** |
| s(year):riverYurappu | 4.251 | 5.227 | 3.66 | **0.0023** |
| s(year):riverAkka | 3.243 | 3.987 | 5.51 | **0.0002** |
| s(year):riverTsugaruishi | 2.883 | 3.581 | 15.60 | **< 0.0001** |
| s(year):riverKatagishi | 4.760 | 5.794 | 4.18 | **0.0009** |
| s(year):riverKitakami | 1.019 | 1.035 | 3.30 | 0.0692 |
| s(year):riverUda | 2.468 | 3.089 | 5.54 | **0.0008** |
| s(year):riverGakko | 2.727 | 3.390 | 6.35 | **0.0002** |
| s(year):riverMiomote | 2.792 | 3.466 | 3.49 | **0.0122** |
| s(year):riverSho | 0.003 | 0.005 | 0.01 | 0.9948 |

edf, estimated degrees of freedom; Ref. df, reference degrees of freedom used in the approximate *F* test for smooth terms.

**Table S10.** Parametric coefficients of the river-specific generalized additive model (GAM) for age-5 egg size after accounting for river effects; *Egg* ~ *s*(year) + *s*(year, by = river) + river (Fig. S7). River coefficients represent differences relative to the reference river (Tokushibetsu). The river-specific GAM for age-5 egg size explained 31.6% of the deviance (adjusted $R^2 = 0.31$).

| Coefficient | Estimate | Standard error | *t* | *p*-value |
|---|---|---|---|---|
| (Intercept) | 0.2491 | 0.0013 | 195.86 | **< 0.0001** |
| riverIshikari | -0.0437 | 0.0020 | -22.05 | **< 0.0001** |
| riverNishibetsu | 0.0024 | 0.0018 | 1.36 | 0.1733 |
| riverTokachi | 0.0369 | 0.0017 | 21.74 | **< 0.0001** |
| riverYurappu | 0.0234 | 0.0017 | 13.87 | **< 0.0001** |
| riverAkka | 0.0102 | 0.0018 | 5.56 | **< 0.0001** |
| riverTsugaruishi | 0.0241 | 0.0017 | 14.28 | **< 0.0001** |
| riverKatagishi | 0.0025 | 0.0018 | 1.40 | 0.1613 |
| riverKitakami | -0.0090 | 0.0020 | -4.56 | **< 0.0001** |
| riverUda | -0.0178 | 0.0020 | -8.73 | **< 0.0001** |
| riverGakko | 0.0007 | 0.0018 | 0.39 | 0.6971 |
| riverMiomote | -0.0051 | 0.0020 | -2.47 | **0.0136** |
| riverSho | -0.0177 | 0.0023 | -7.81 | **< 0.0001** |

**Table S11.** Approximate significance of smooth terms in the river-specific generalized additive model (GAM) for age-5 egg size after accounting for river effects; *Egg* ~ *s*(year) + *s*(year, by = river) + river (Fig. 3). Non-significant river-specific smooth terms indicate that temporal trends in those rivers did not differ significantly from the overall smooth trend.

| Smooth term | edf | Ref. df | *F* | *p*-value |
|---|---|---|---|---|
| s(year) | 8.641 | 8.933 | 8.06 | **< 0.0001** |
| s(year):riverTokushibetsu | 3.529 | 4.372 | 5.09 | **0.0003** |
| s(year):riverIshikari | 5.164 | 6.221 | 2.08 | 0.0687 |
| s(year):riverNishibetsu | 6.355 | 7.476 | 4.77 | **< 0.0001** |
| s(year):riverTokachi | 7.516 | 8.429 | 3.07 | **0.0137** |
| s(year):riverYurappu | 1.000 | 1.001 | 1.28 | 0.2582 |
| s(year):riverAkka | 2.462 | 3.060 | 2.42 | 0.0635 |
| s(year):riverTsugaruishi | 2.785 | 3.465 | 3.45 | **0.0113** |
| s(year):riverKatagishi | 2.153 | 2.699 | 1.39 | 0.1892 |
| s(year):riverKitakami | 1.000 | 1.000 | 1.51 | 0.2188 |
| s(year):riverUda | 1.056 | 1.108 | 1.67 | 0.1791 |
| s(year):riverGakko | 5.319 | 6.395 | 1.76 | 0.0823 |
| s(year):riverMiomote | 1.712 | 2.136 | 1.38 | 0.2516 |
| s(year):riverSho | 3.283 | 4.269 | 1.35 | 0.1760 |

edf, estimated degrees of freedom; Ref. df, reference degrees of freedom used in the approximate *F* test for smooth terms.

**Table S12.** Parametric coefficients of the hierarchical generalized additive model (GAM) for female fork length (FL) after accounting for age, and river effects; $FL \sim s(\text{year}) + s(\text{year, by = river}) + \text{age} + \text{river}$. River coefficients represent differences relative to the reference river (Tokushibetsu). The hierarchical GAM for female FL explained 43.7% of the deviance (adjusted $R^2 = 0.44$).

| Coefficient | Estimate | Standard error | *t* | *p*-value |
|---|---|---|---|---|
| (Intercept) | 51.3089 | 0.2368 | 216.68 | < **0.0001** |
| age | 3.6330 | 0.0496 | 73.31 | < **0.0001** |
| riverIshikari | -0.4849 | 0.1224 | -3.96 | < **0.0001** |
| riverNishibetsu | -1.3463 | 0.1193 | -11.29 | < **0.0001** |
| riverTokachi | 0.3356 | 0.1179 | 2.85 | **0.0044** |
| riverYurappu | 3.4243 | 0.1191 | 28.76 | < **0.0001** |
| riverAkka | 0.3545 | 0.1244 | 2.85 | **0.0044** |
| riverTsugaruishi | 5.1610 | 0.1204 | 42.87 | < **0.0001** |
| riverKatagishi | 0.9050 | 0.1214 | 7.45 | < **0.0001** |
| riverKitakami | 0.5462 | 0.1265 | 4.32 | < **0.0001** |
| riverUda | 0.5693 | 0.1309 | 4.35 | < **0.0001** |
| riverGakko | 4.9946 | 0.1212 | 41.20 | < **0.0001** |
| riverMiomote | 3.8414 | 0.1259 | 30.50 | < **0.0001** |
| riverSho | 2.4251 | 0.1285 | 18.88 | < **0.0001** |

**Table S13.** Approximate significance of smooth terms in the hierarchical generalized additive model (GAM) for female fork length (FL) after accounting for age, and river effects; *FL* ~ *s*(year) + *s*(year, by = river) + age + river. Non-significant river-specific smooth terms indicate that temporal trends in those rivers did not differ significantly from the overall smooth trend.

| Smooth term | edf | Ref. df | *F* | *p*-value |
|---|---|---|---|---|
| s(year) | 8.906 | 8.981 | 60.12 | < **0.0001** |
| s(year):riverTokushibetsu | 2.476 | 3.084 | 1.67 | 0.1675 |
| s(year):riverIshikari | 5.742 | 6.875 | 3.47 | **0.0016** |
| s(year):riverNishibetsu | 8.676 | 8.963 | 13.09 | < **0.0001** |
| s(year):riverTokachi | 3.662 | 4.528 | 1.59 | 0.1531 |
| s(year):riverYurappu | 5.890 | 7.029 | 3.64 | **0.0006** |
| s(year):riverAkka | 3.606 | 4.447 | 5.13 | **0.0002** |
| s(year):riverTsugaruishi | 3.725 | 4.605 | 4.03 | **0.0018** |
| s(year):riverKatagishi | 5.945 | 7.094 | 4.47 | < **0.0001** |
| s(year):riverKitakami | 1.024 | 1.035 | 0.58 | 0.4536 |
| s(year):riverUda | 3.034 | 3.777 | 2.13 | 0.0874 |
| s(year):riverGakko | 7.191 | 8.205 | 5.62 | < **0.0001** |
| s(year):riverMiomote | 6.013 | 7.063 | 4.04 | **0.0003** |
| s(year):riverSho | 5.906 | 7.033 | 3.18 | **0.0022** |

edf, estimated degrees of freedom; Ref. df, reference degrees of freedom used in the approximate *F* test for smooth terms.

**Table S14.** Parametric coefficients of the hierarchical generalized additive model (GAM) for egg size after accounting for female fork length (FL), age, and river effects; *Egg* ~ *s*(year) + *s*(year, by = river) + *FL* + age + river. River coefficients represent differences relative to the reference river (Tokushibetsu). The hierarchical GAM for egg size explained 41.9% of the deviance (adjusted $R^2$ = 0.42).

| Coefficient | Estimate | Standard error | *t* | *p*-value |
|---|---|---|---|---|
| (Intercept) | 0.0298 | 0.0036 | 8.29 | **< 0.0001** |
| FL | 0.0028 | 0.0001 | 47.88 | **< 0.0001** |
| age | 0.0053 | 0.0005 | 11.28 | **< 0.0001** |
| riverIshikari | -0.0373 | 0.0010 | -35.73 | **< 0.0001** |
| riverNishibetsu | 0.0082 | 0.0010 | 7.99 | **< 0.0001** |
| riverTokachi | 0.0343 | 0.0010 | 34.01 | **< 0.0001** |
| riverYurappu | 0.0128 | 0.0010 | 12.42 | **< 0.0001** |
| riverAkka | 0.0076 | 0.0011 | 7.12 | **< 0.0001** |
| riverTsugaruishi | 0.0072 | 0.0011 | 6.78 | **< 0.0001** |
| riverKatagishi | 0.0014 | 0.0010 | 1.39 | 0.1640 |
| riverKitakami | -0.0096 | 0.0011 | -8.86 | **< 0.0001** |
| riverUda | -0.0195 | 0.0011 | -17.67 | **< 0.0001** |
| riverGakko | -0.0131 | 0.0011 | -12.19 | **< 0.0001** |
| riverMiomote | -0.0131 | 0.0011 | -12.01 | **< 0.0001** |
| riverSho | -0.0220 | 0.0011 | -19.98 | **< 0.0001** |

**Table S15.** Approximate significance of smooth terms in the hierarchical generalized additive model (GAM) for egg size after accounting for female fork length (FL), age, and river effects; *Egg* ~ *s*(year) + *s*(year, by = river) + *FL* + age + river. Non-significant river-specific smooth terms indicate that temporal trends in those rivers did not differ significantly from the overall smooth trend.

| Smooth term | edf | Ref. df | *F* | *p*-value |
|---|---|---|---|---|
| s(year) | 8.427 | 8.853 | 17.48 | **< 0.0001** |
| s(year):riverTokushibetsu | 7.715 | 8.555 | 14.20 | **< 0.0001** |
| s(year):riverIshikari | 6.065 | 7.205 | 2.40 | **0.0166** |
| s(year):riverNishibetsu | 6.341 | 7.472 | 5.93 | **< 0.0001** |
| s(year):riverTokachi | 8.516 | 8.924 | 8.31 | **< 0.0001** |
| s(year):riverYurappu | 1.009 | 1.017 | 1.70 | 0.1917 |
| s(year):riverAkka | 4.921 | 5.984 | 1.26 | 0.2394 |
| s(year):riverTsugaruishi | 2.429 | 3.029 | 4.20 | **0.0054** |
| s(year):riverKatagishi | 1.024 | 1.048 | 1.86 | 0.1659 |
| s(year):riverKitakami | 1.002 | 1.003 | 0.17 | 0.6852 |
| s(year):riverUda | 1.011 | 1.022 | 0.08 | 0.7962 |
| s(year):riverGakko | 7.963 | 8.704 | 4.65 | **< 0.0001** |
| s(year):riverMiomote | 0.006 | 0.012 | 0.00 | 0.9957 |
| s(year):riverSho | 5.360 | 6.463 | 9.28 | **< 0.0001** |

edf, estimated degrees of freedom; Ref. df, reference degrees of freedom used in the approximate *F* test for smooth terms.

**Table S16.** Comparison of exponential models without a breakpoint and segmented regression models for temporal trends in egg size, return rate, and abundance of Japanese chum salmon. Models were compared using Akaike's information criterion (AIC). ΔAIC was calculated as AIC(no breakpoint) − AIC(segmented).

| Model | Age-4 egg size | Age-5 egg size | Return rate | Abundance |
|---|---|---|---|---|
| No breakpoint | -94.08 | -91.27 | 10.99 | 17.02 |
| Segmented | -94.22 | -89.62 | 10.66 | **4.79** |
| ΔAIC | 0.14 | -1.65 | 0.33 | **12.23** |
| Breakpoint | 2003.0 | 2018.0 | 2017.8 | 2005.3 |

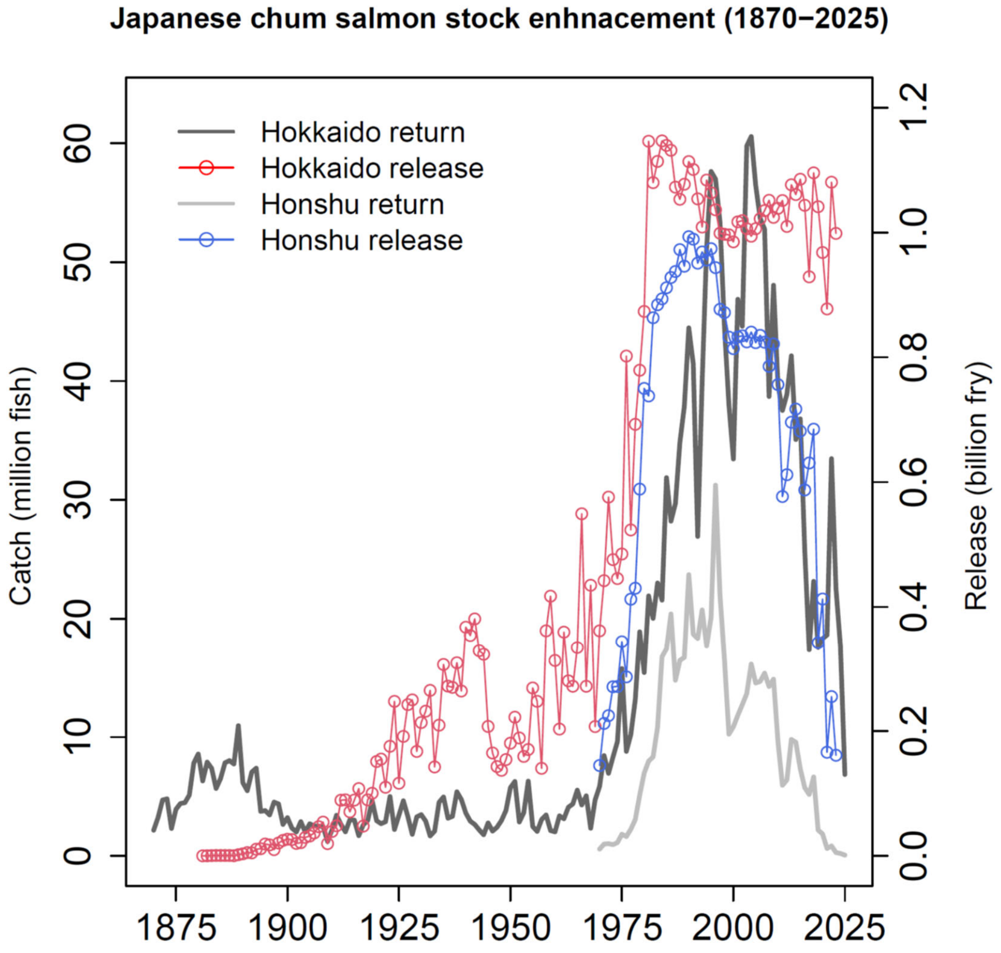


**Fig. S1.** Historical trends in catch and release of Japanese chum salmon over the past 150 years (1870–2025). Updated from Kitada (2020). The Japanese chum salmon enhancement program has released approximately 102 billion fry between 1881 and 2023. Returns of Japanese chum salmon increased markedly following the rapid expansion of hatchery releases after 1975, peaking in 1996. However, after a regional peak in Hokkaido in 2004, returns fluctuated but declined overall, falling below 10 million fish in 2025 and below the maximum level observed prior to hatchery expansion.

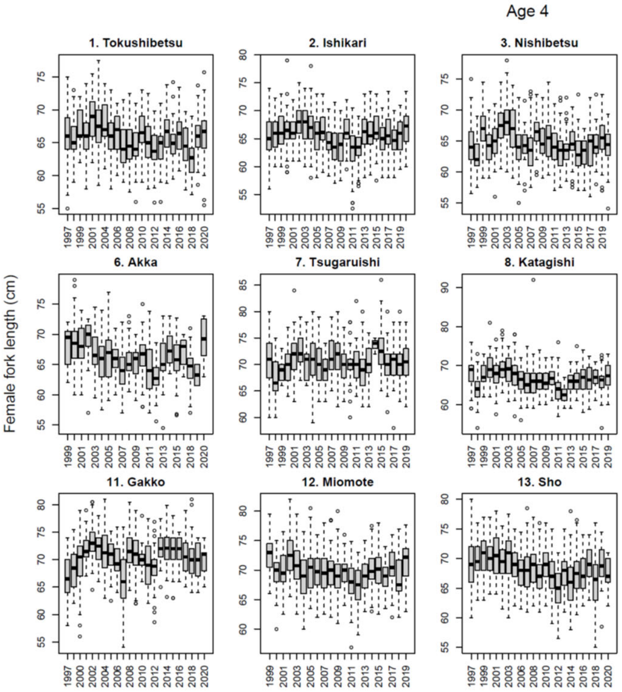


**Fig. S2.** Boxplots of female fork length (FL) of age-4 chum salmon in 13 rivers in Japan from 1997 to 2020 brood year (sample sizes are given in Tables S1, S2). Boxplots show the distribution of individual observations by year for each river, illustrating substantial interannual and among-river variability underlying the annual means used in the GAM analyses. The boxplots show the data before the exclusion of extreme values. Extreme values were excluded from the analyses because their validity could not be confirmed.

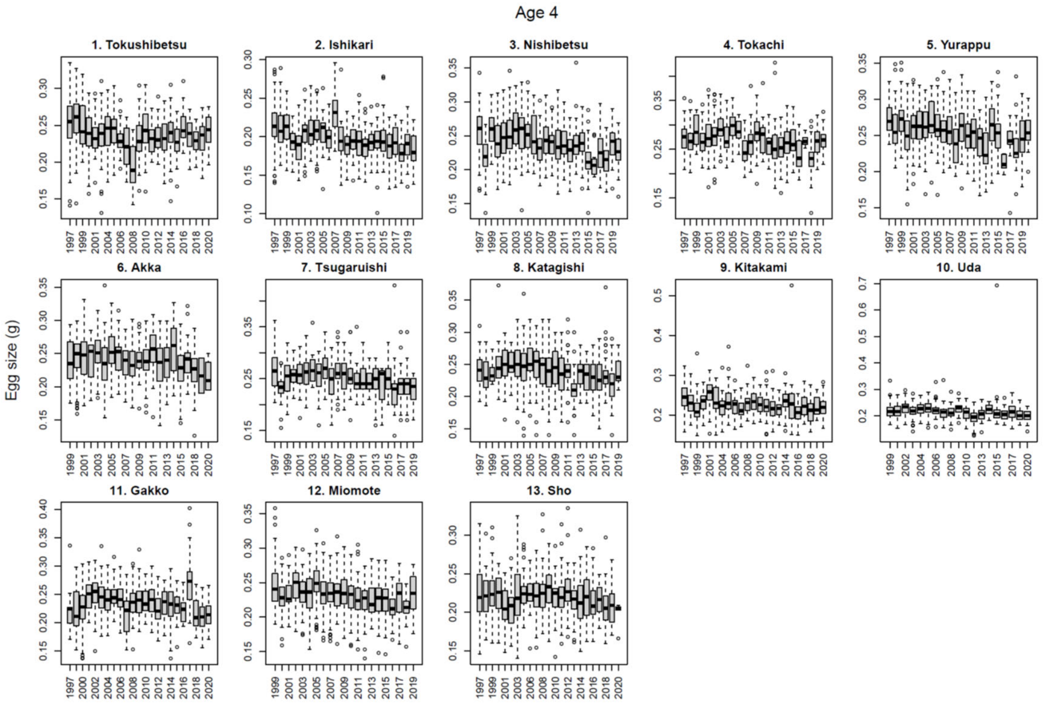


**Fig. S3.** Boxplots of egg size of age-4 chum salmon in 13 rivers in Japan from 1997 to 2020 brood year (sample sizes are given in Tables S1, S2). Boxplots show the distribution of individual observations by year for each river, illustrating substantial interannual and among-river variability underlying the annual means used in the GAM analyses. The boxplots show the data before the exclusion of extreme values. Extreme values were excluded from the analyses because their validity could not be confirmed.

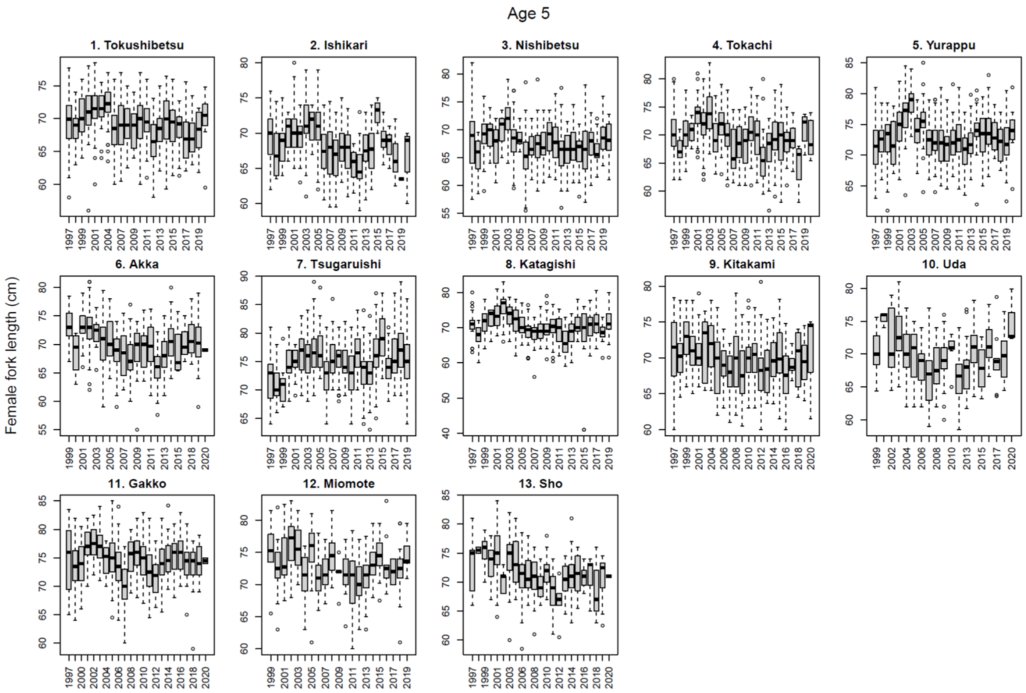


**Fig. S4.** Boxplots of female fork length (FL) of age-5 chum salmon in 13 rivers in Japan from 1997 to 2020 brood year (sample sizes are given in Tables S1, S3). Boxplots show the distribution of individual observations by year for each river, illustrating substantial interannual and among-river variability underlying the annual means used in the GAM analyses. The boxplots show the data before the exclusion of extreme values. Extreme values were excluded from the analyses because their validity could not be confirmed.

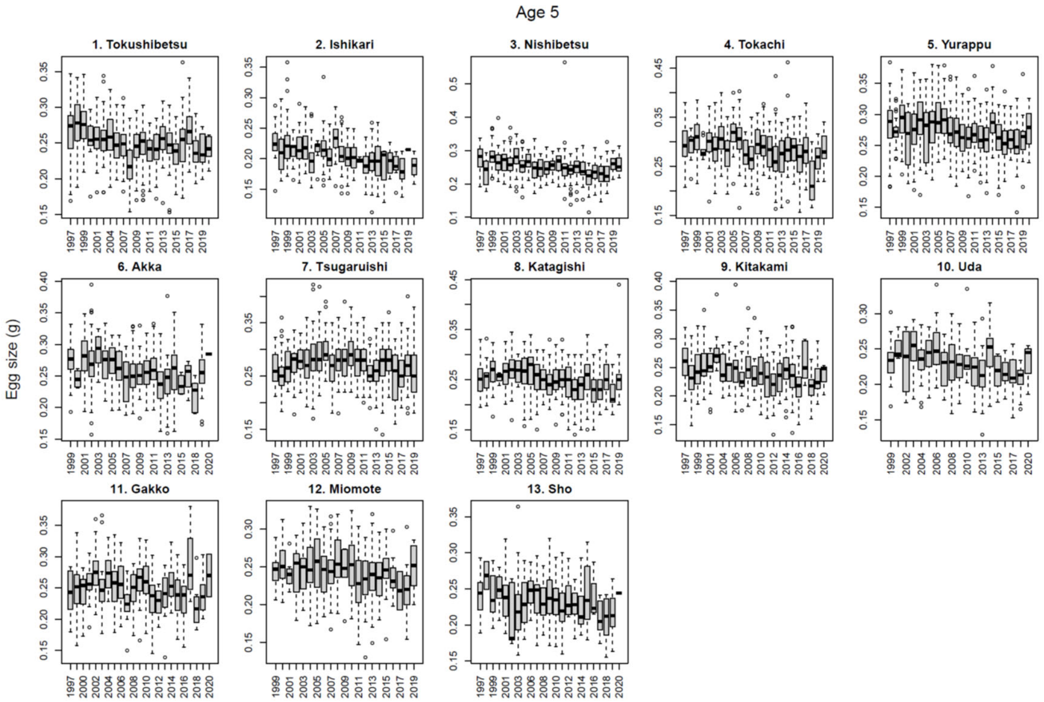


**Fig. S5.** Boxplots of egg size of age-5 chum salmon in 13 rivers in Japan from 1997 to 2020 brood year (sample sizes are given in Tables S1, S3). Boxplots show the distribution of individual observations by year for each river, illustrating substantial interannual and among-river variability underlying the annual means used in the GAM analyses. Extreme values were retained because their validity could not be confirmed.

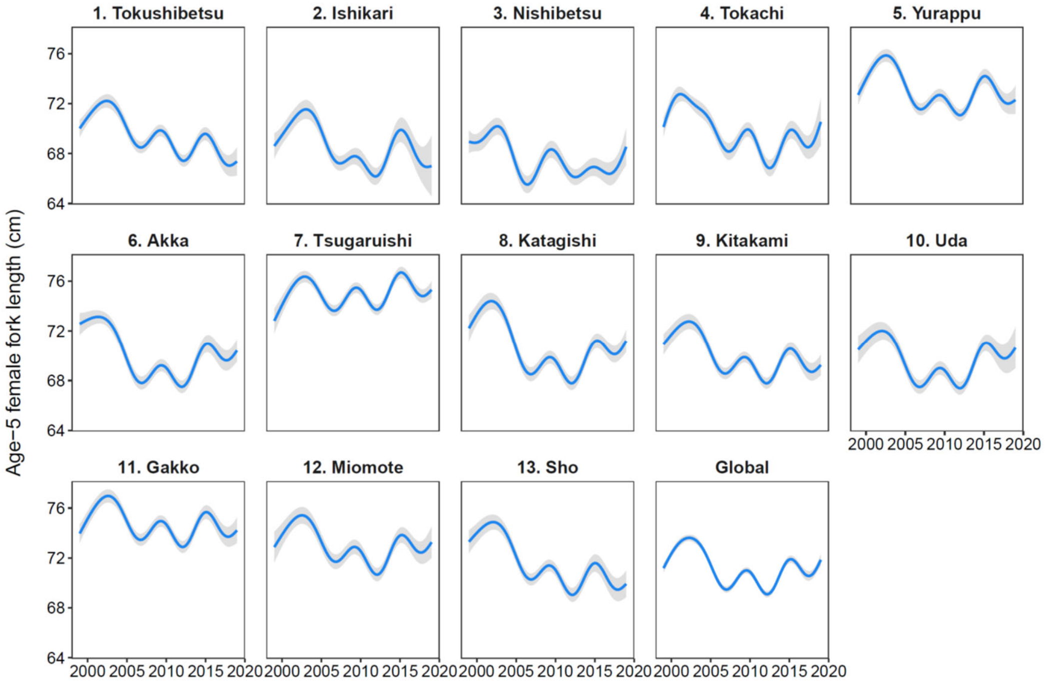


**Fig. S6.** Predicted temporal trends in female fork length (FL) for age-5 chum salmon estimated from generalized additive models fitted to individual-level data from 8,862 females sampled from 13 hatchery-enhanced rivers for the 1999–2019 brood years. Blue lines show the fitted GAM smooths, and gray shaded areas indicate the 95% confidence intervals. The global smooth represents the common temporal trend estimated from a separate GAM with a single common smooth of year fitted to all observations pooled across the 13 rivers.

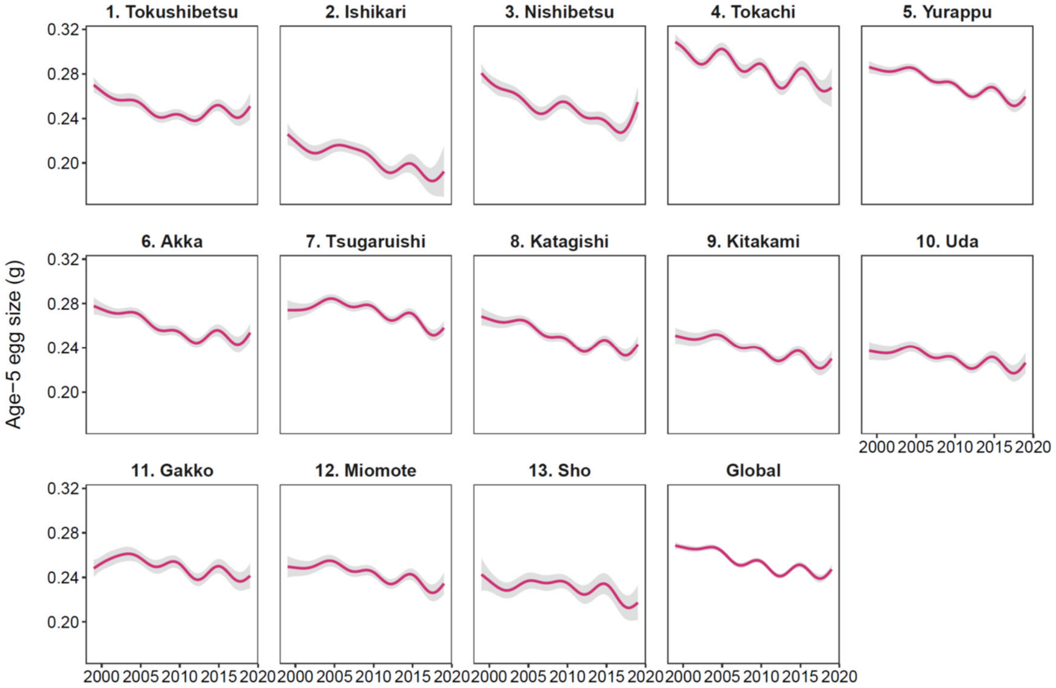


**Fig. S7.** Predicted temporal trends in egg size for age-5 chum salmon estimated from generalized additive models fitted to individual-level data from 8,862 females sampled from 13 hatchery-enhanced rivers for the 1999–2019 brood years. Pink lines show the fitted GAM smooths, and gray shaded areas indicate the 95% confidence intervals. The global smooth represents the common temporal trend estimated from a separate GAM with a single common smooth of year fitted to all observations pooled across the 13 rivers.

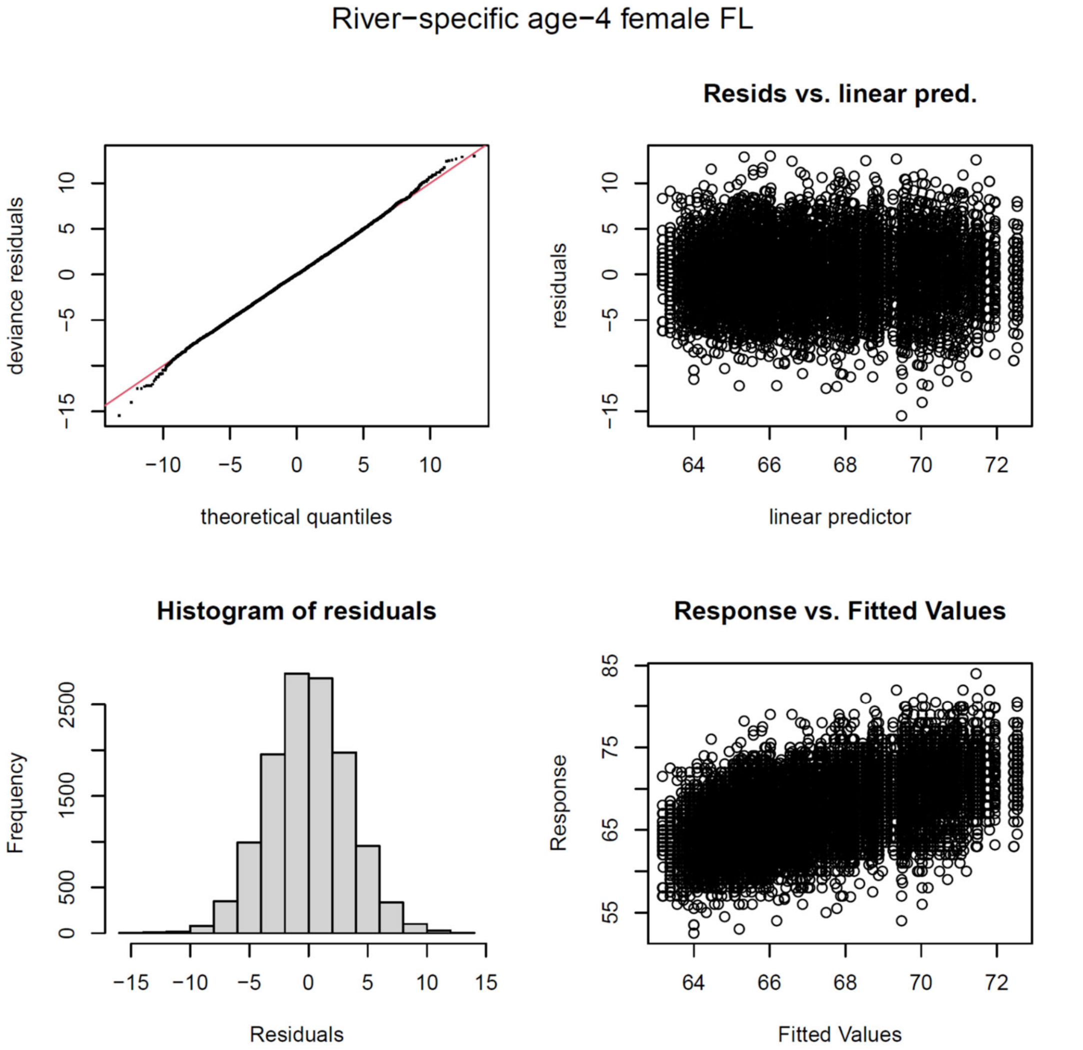


**Fig. S8.** Diagnostic plots for the river-specific generalized additive model (GAM) fitted to individual-level female fork length (FL) data from 12,428 age-4 female chum salmon collected from 13 hatchery-enhanced rivers. The panels show (upper left) the normal Q–Q plot of deviance residuals, (upper right) residuals versus linear predictor, (lower left) the histogram of residuals, and (lower right) observed versus fitted values. The diagnostic plots indicate no substantial departures from model assumptions.

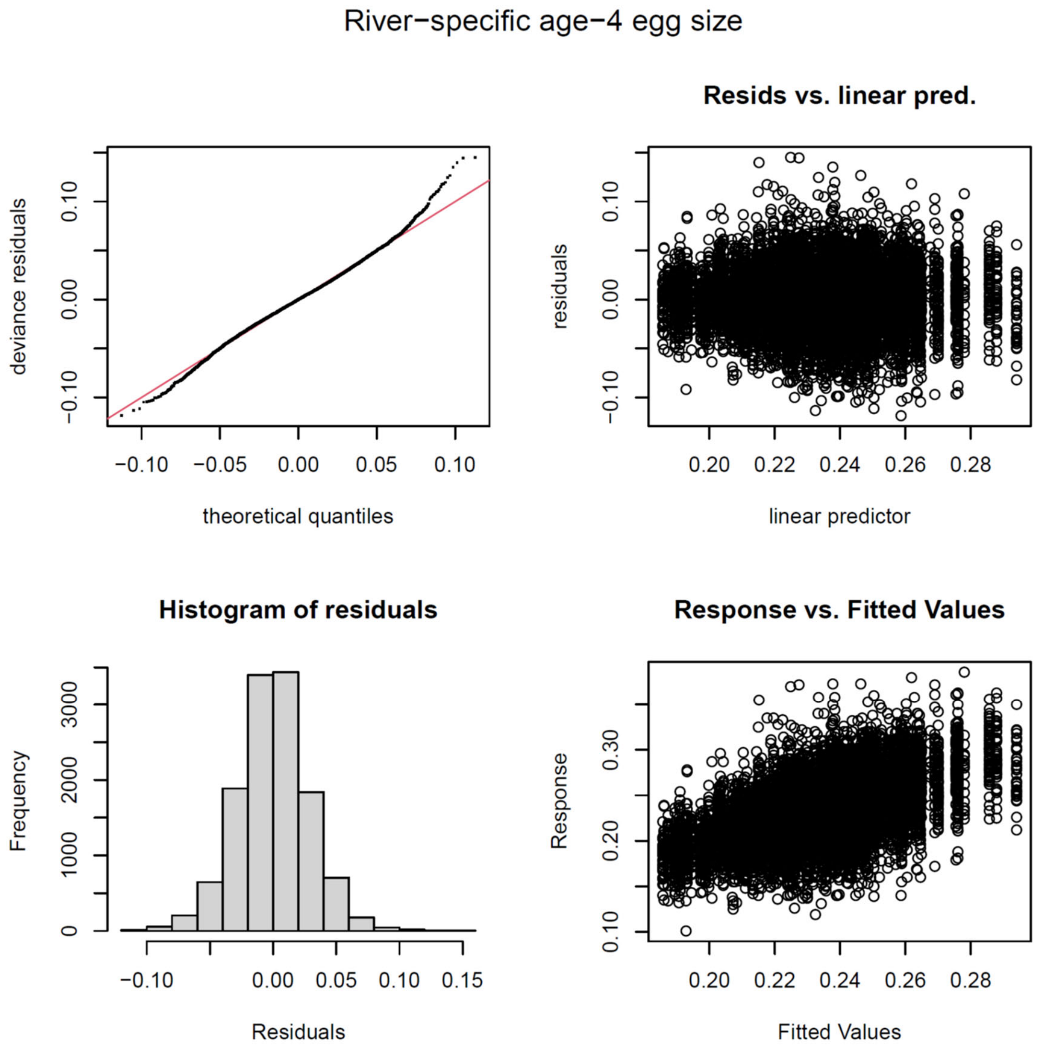


**Fig. S9** Diagnostic plots for the river-specific generalized additive model (GAM) fitted to individual-level egg size data from 12,428 age-4 female chum salmon collected from 13 hatchery-enhanced rivers. The panels show (upper left) the normal Q–Q plot of deviance residuals, (upper right) residuals versus linear predictor, (lower left) the histogram of residuals, and (lower right) observed versus fitted values. The diagnostic plots indicate no substantial departures from model assumptions.

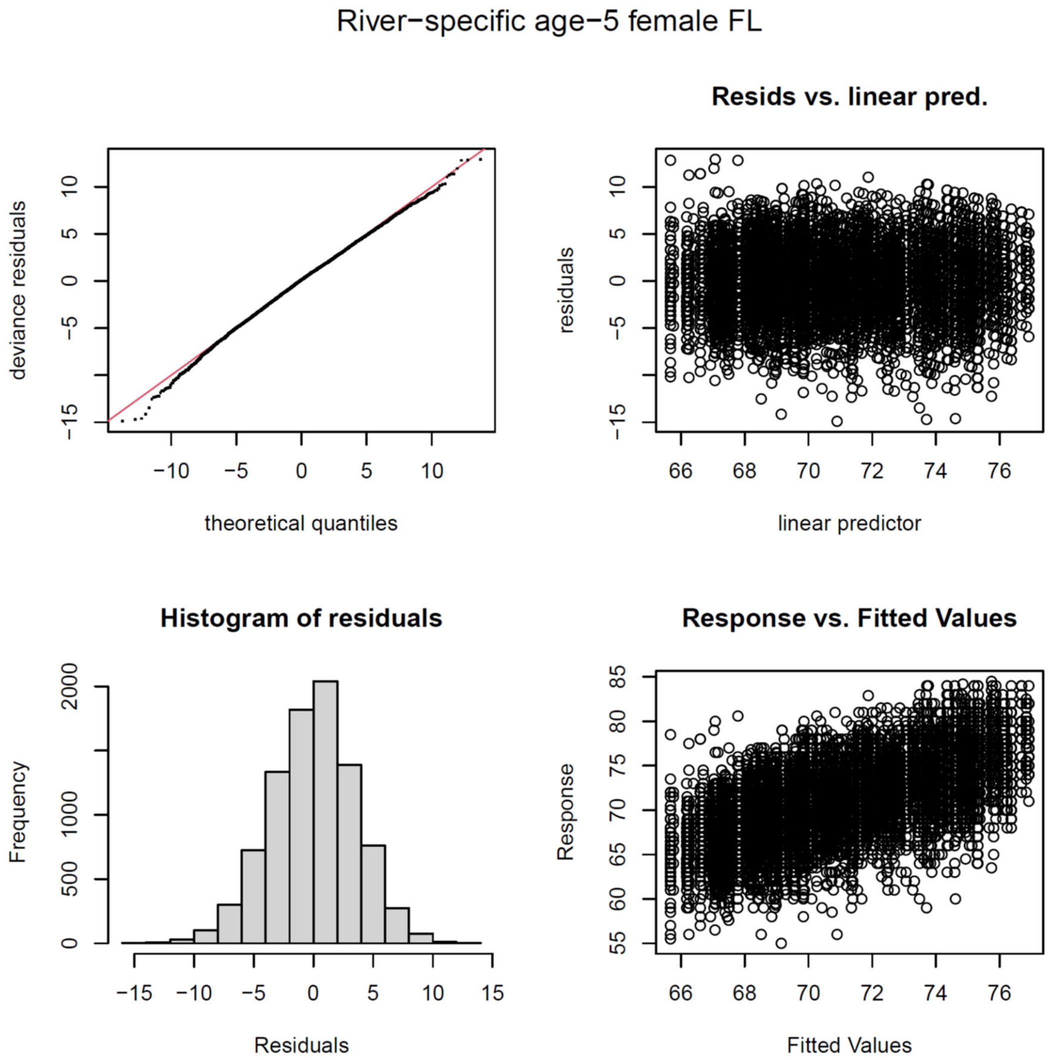


**Fig. S10.** Diagnostic plots for the river-specific generalized additive model (GAM) fitted to individual-level female fork length (FL) data from 8,862 age-5 female chum salmon collected from 13 hatchery-enhanced rivers. The panels show (upper left) the normal Q–Q plot of deviance residuals, (upper right) residuals versus linear predictor, (lower left) the histogram of residuals, and (lower right) observed versus fitted values. The diagnostic plots indicate no substantial departures from model assumptions.

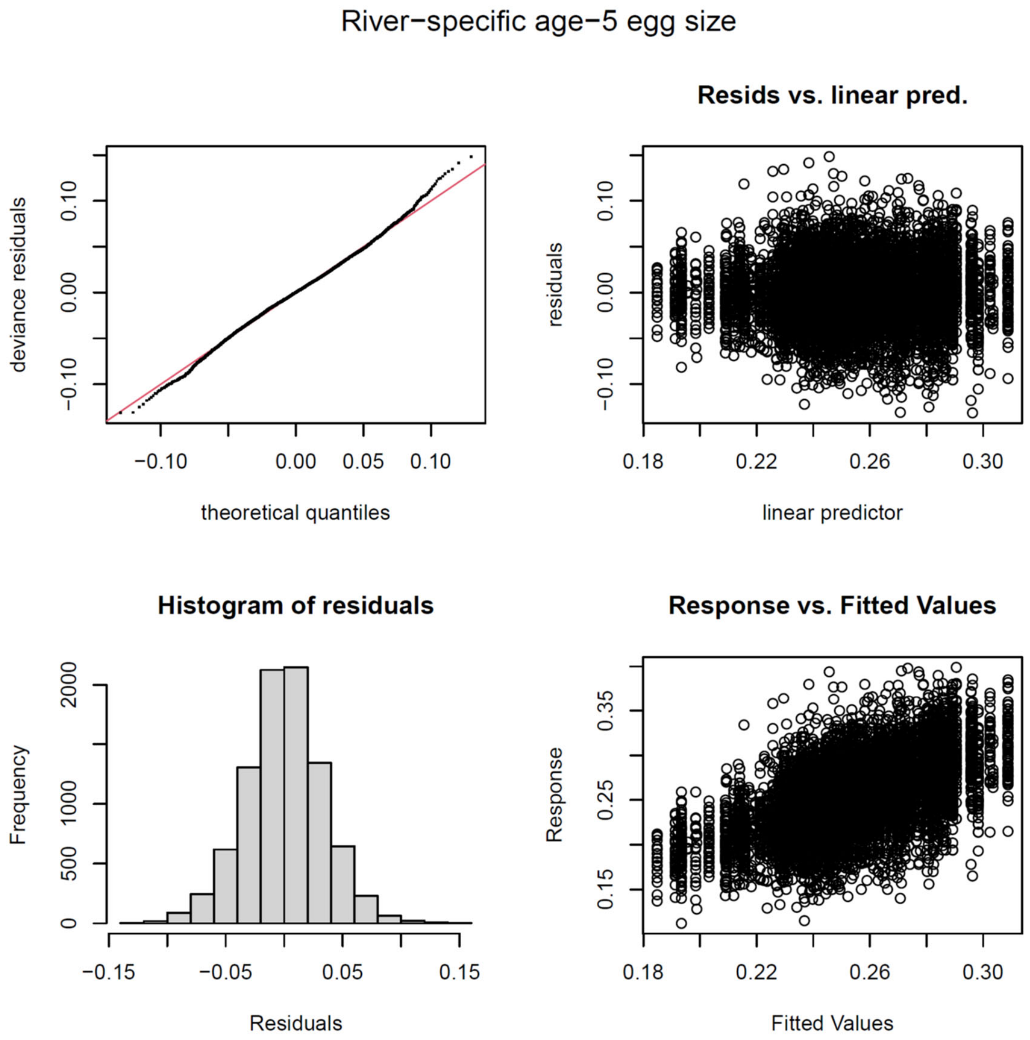


**Fig. S11.** Diagnostic plots for the river-specific generalized additive model (GAM) fitted to individual-level egg size data from 8,862 age-5 female chum salmon collected from 13 hatchery-enhanced rivers. The panels show (upper left) the normal Q–Q plot of deviance residuals, (upper right) residuals versus linear predictor, (lower left) the histogram of residuals, and (lower right) observed versus fitted values. The diagnostic plots indicate no substantial departures from model assumptions.

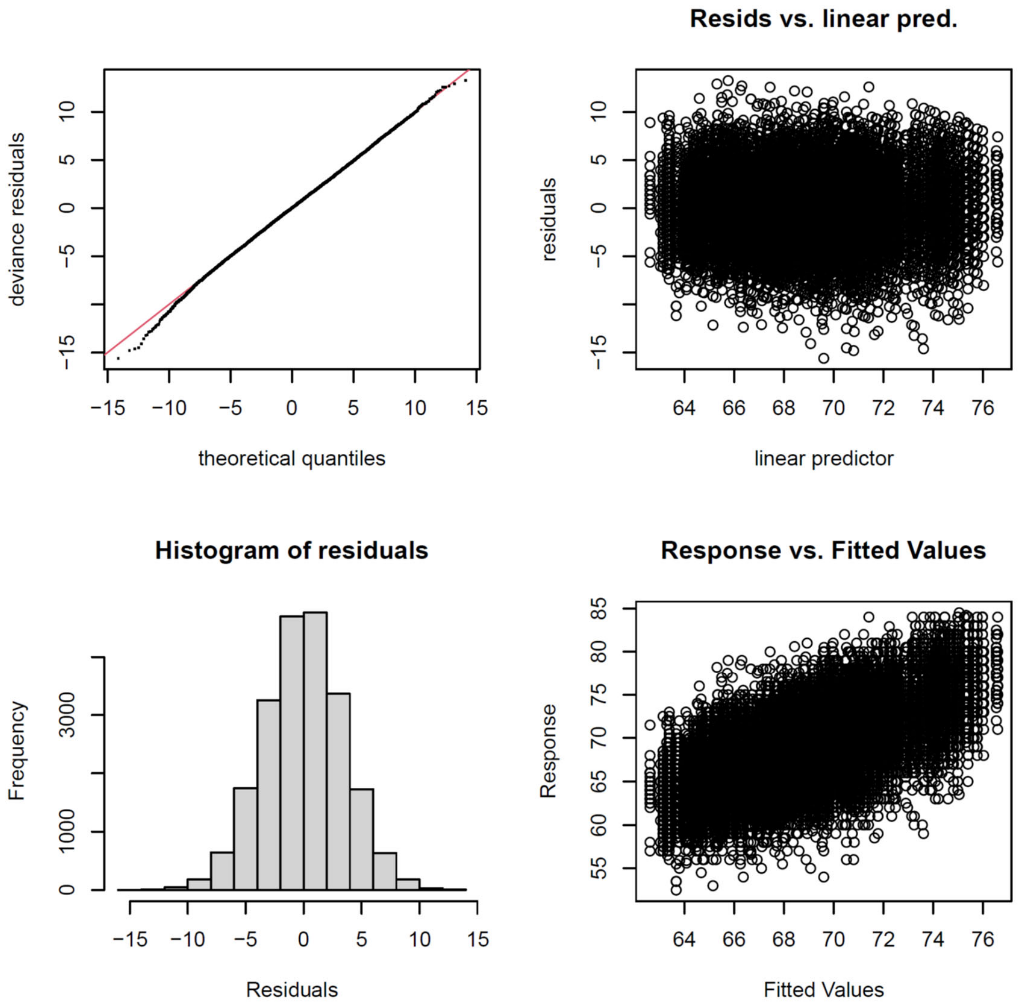


**Fig. S12.** Diagnostic plots for the nationwide hierarchical generalized additive model (GAM) fitted to individual-level female fork length (FL) data from 21,290 age-4 and age-5 female chum salmon collected from 13 hatchery-enhanced rivers. The panels show (upper left) the normal Q–Q plot of deviance residuals, (upper right) residuals versus linear predictor, (lower left) the histogram of residuals, and (lower right) observed versus fitted values. The diagnostic plots indicate no substantial departures from model assumptions.

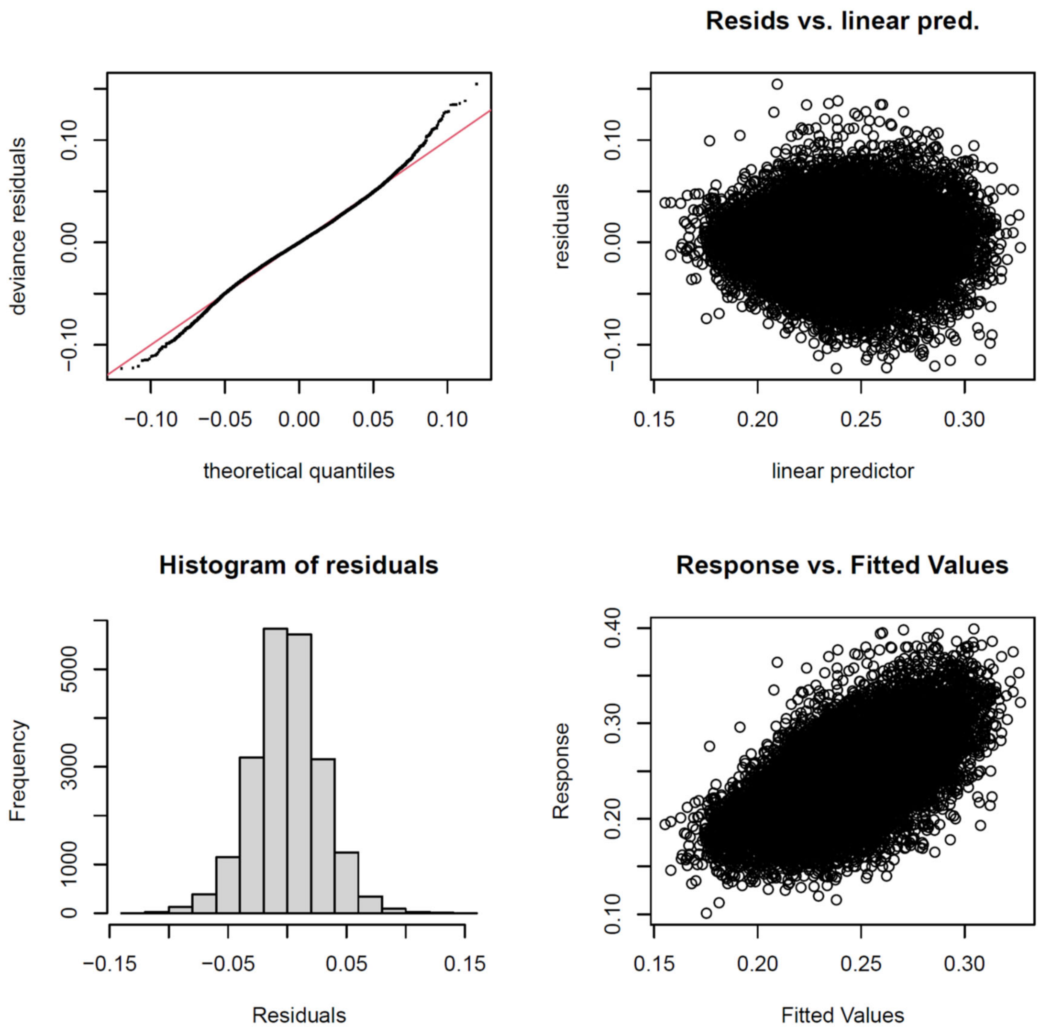


**Fig. S13.** Diagnostic plots for the nationwide hierarchical generalized additive model (GAM) fitted to individual-level egg-size data from 21,290 age-4 and age-5 female chum salmon collected from 13 hatchery-enhanced rivers. The panels show (upper left) the normal Q–Q plot of deviance residuals, (upper right) deviance residuals versus linear predictor, (lower left) the histogram of residuals, and (lower right) observed versus fitted values. The diagnostic plots indicate no substantial departures from model assumptions.

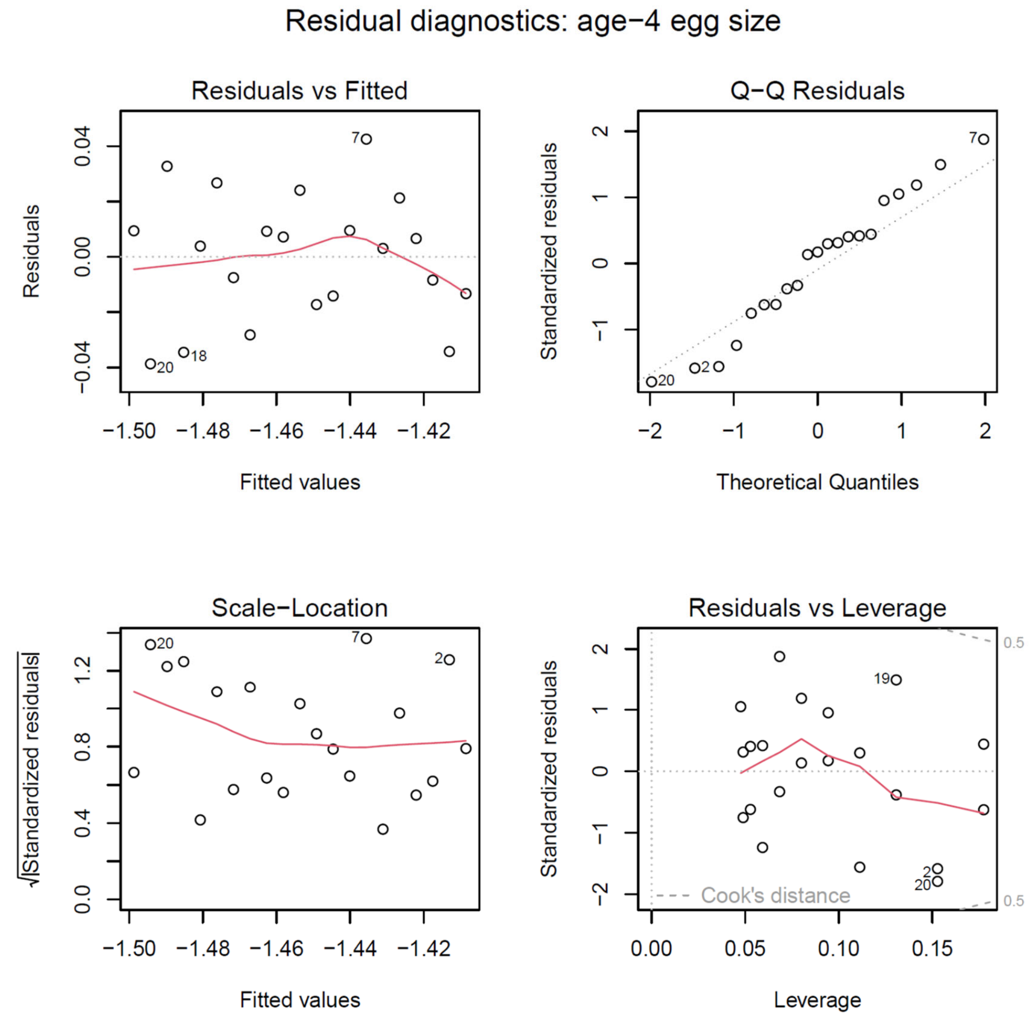


**Fig. S14.** Diagnostic plots for the exponential decay model fitted to annual mean age-4 egg size of Japanese chum salmon (Fig. 6A). The model was used to estimate the long-term decline rate of egg size. The four panels show residuals versus fitted values, normal Q–Q plots, scale-location plots, and residuals versus leverage plots for assessing model assumptions and influential observations.

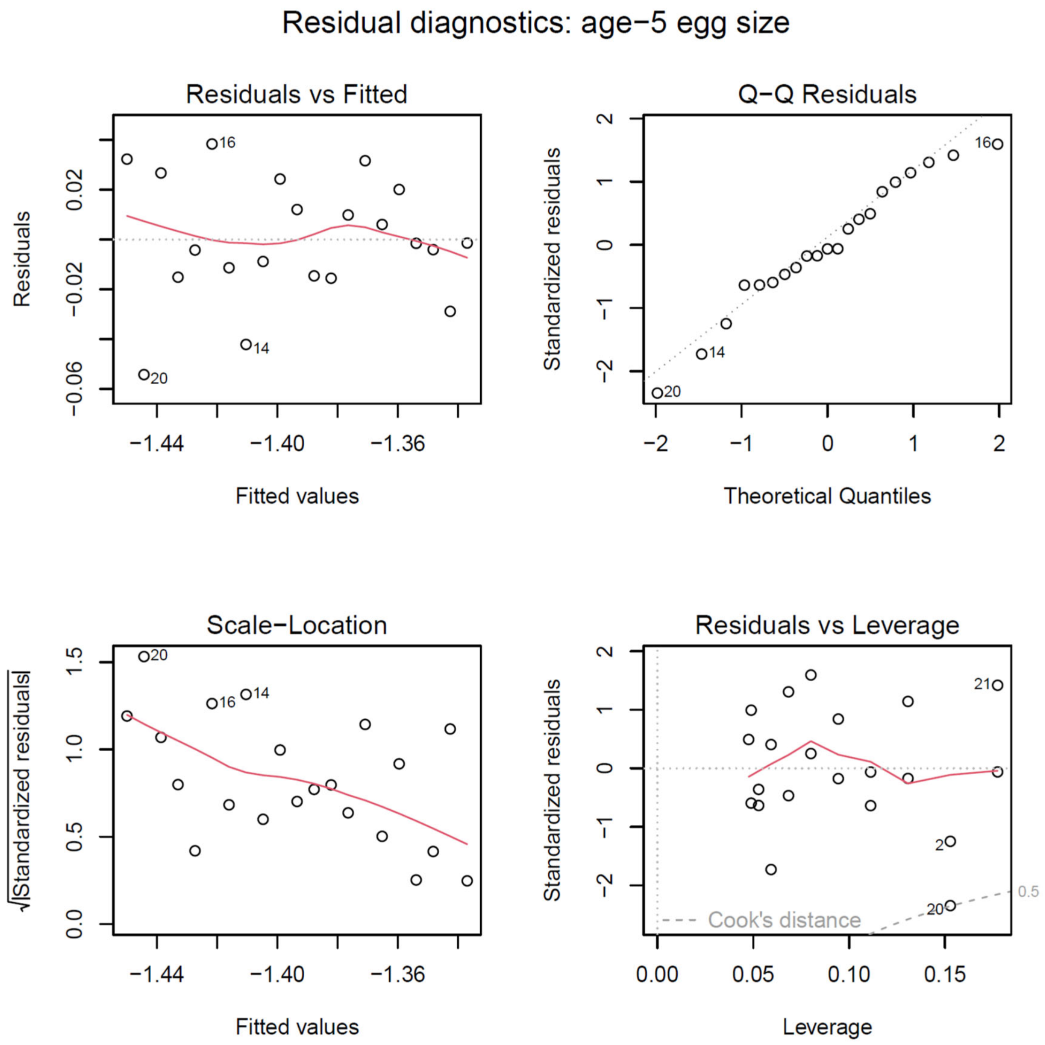


**Fig. S15.** Diagnostic plots for the exponential decay model fitted to annual mean age-5 egg size of Japanese chum salmon (Fig. 6B). The model was used to estimate the long-term decline rate of egg size. The four panels show residuals versus fitted values, normal Q–Q plots, scale-location plots, and residuals versus leverage plots for assessing model assumptions and influential observations.

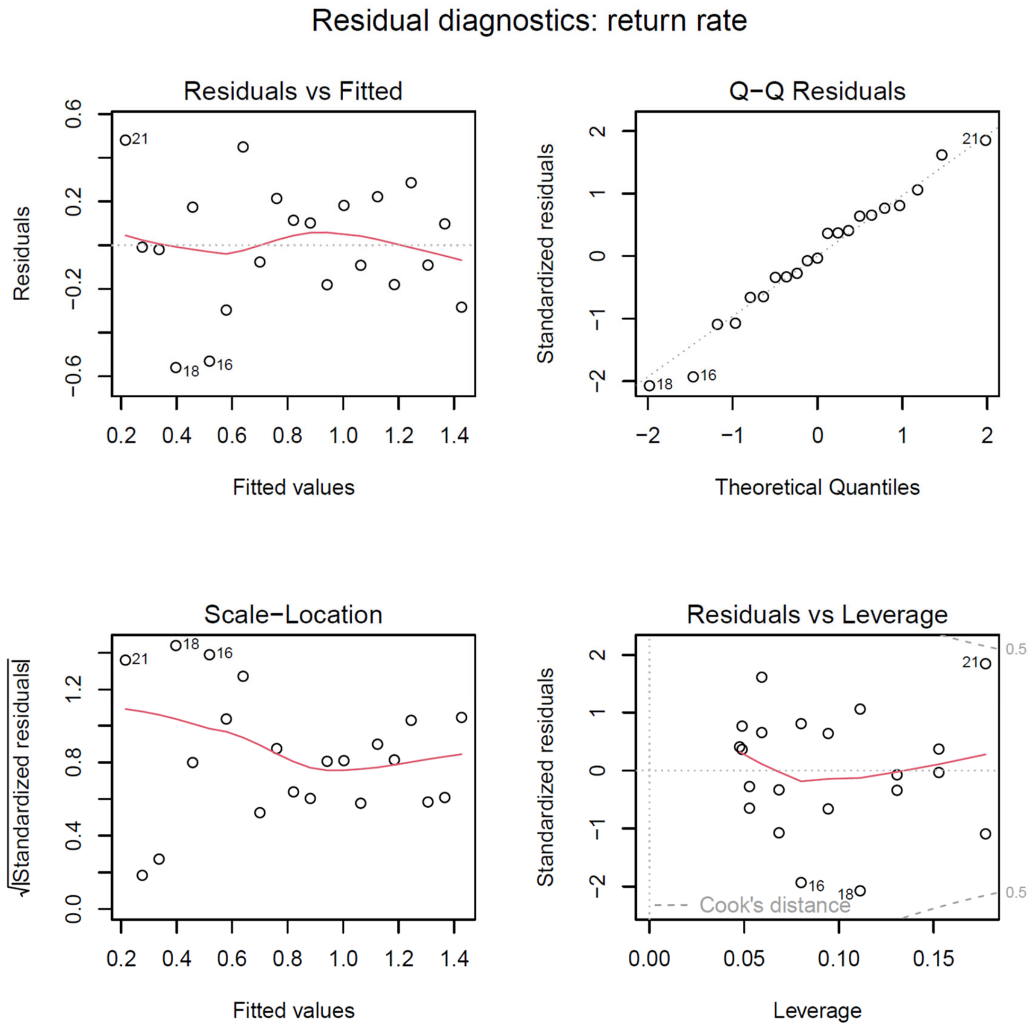


**Fig. S16.** Diagnostic plots for the exponential decay model fitted to brood-year-specific return rates of Japanese chum salmon (Fig. 6C). The model was used to estimate the long-term decline rate of return rate. The four panels show residuals versus fitted values, normal Q–Q plots, scale-location plots, and residuals versus leverage plots for assessing model assumptions and influential observations.

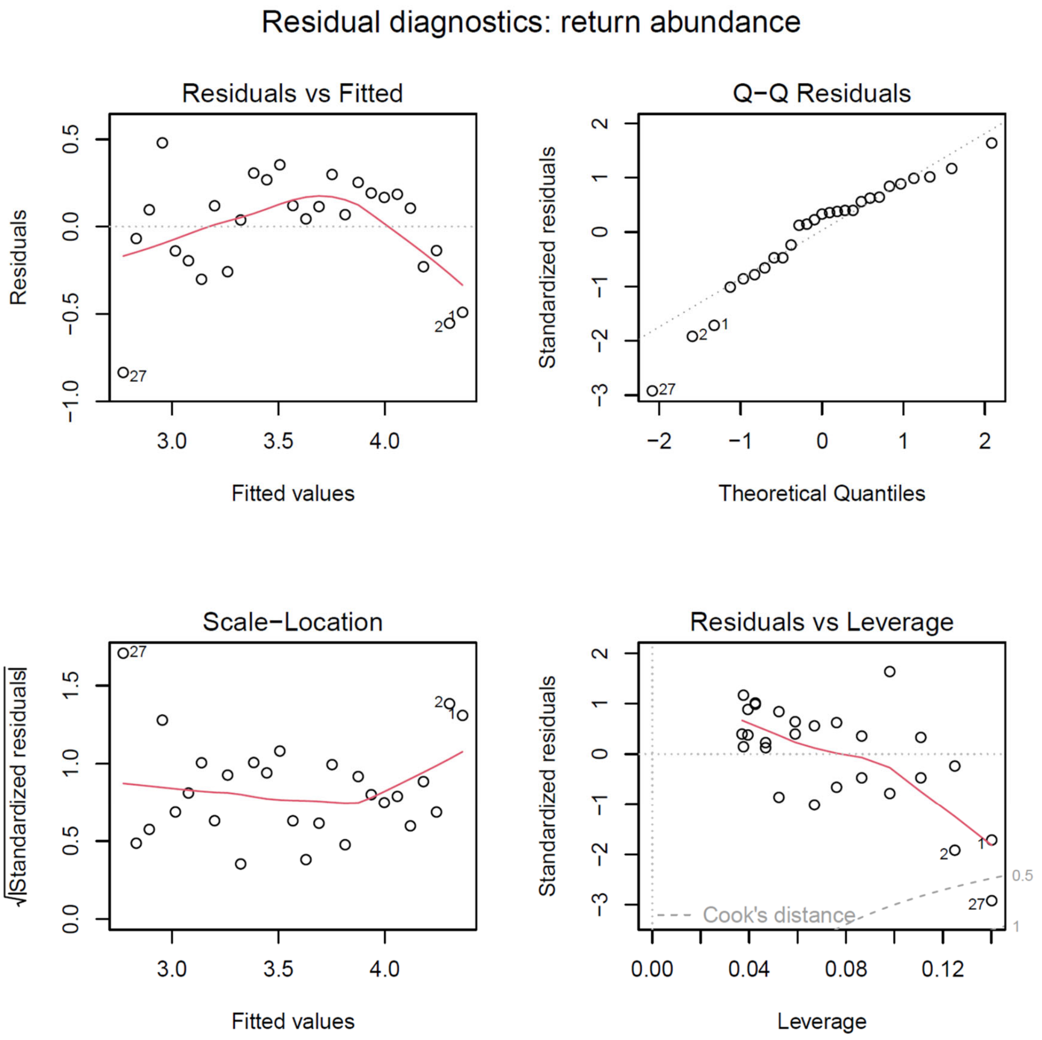


**Fig. S17.** Diagnostic plots for the exponential decay model fitted to annual return abundance of Japanese chum salmon (Fig. 6D). The model was used to estimate the long-term decline rate of abundance. The four panels show residuals versus fitted values, normal Q–Q plots, scale-location plots, and residuals versus leverage plots for assessing model assumptions and influential observations.
